# Identifying molecular instructions to hard-wire a sensory neuron’s synaptic connectivity

**DOI:** 10.64898/2026.04.22.720075

**Authors:** Júnia Vieira dos Santos, Renee Yin Yu, Andrea Araceli Terceros, Peter Mussells Pires, Denisa Rusu, Vedrana Cvetkovska, Alyeri Bucio-Méndez, Tsung-Jung Lin, Farida Emran, Haig Djambazian, Pierre Bérubé, Robert Sladek, Brian E. Chen

**Affiliations:** Centre for Research in Neuroscience, Research Institute of the McGill University Health Centre, Montréal, Québec, Canada, H3G 1A4; Génome Québec Innovation Centre, Montréal, Québec, Canada, H3A 0G1; Departments of Medicine and Neurology & Neurosurgery, McGill University, Montréal, Québec, Canada, H3G 1A4

## Abstract

The neural circuitry underlying innate behaviours must be pre-specified using precise molecular instructions. However, it is not clear what specific molecules are required to generate one neuron’s wiring pattern that make it distinct from a neighboring neuron’s wiring pattern. Here, we repeatedly sequenced three identified *Drosophila* sensory neurons that each have a unique and stereotyped hard-wired synaptic connectivity, to determine the cell-surface molecules that distinguish their identities. Repeated sequencing of the same neuron across different animals revealed that the variability of transcription is < 1%. We find that fewer than 100 cell-surface molecules can distinguish a mechanosensory neuron from a chemosensory neuron. Functional characterization of the cell-surface receptors verified their different roles in axonal and synaptic targeting. Finally, we expressed combinations of the different cell-surface receptors to mis-wire the chemosensory neuron and increase its axonal branching.

## Introduction

Critical behaviours that are required for an animal’s survival are hard-wired into the genome. The neural circuitry underlying these innate behaviours must be pre-specified using precise molecular instructions so that they wire up properly without experience-dependent refinement. Previous research has uncovered how specific molecules can guide, attract, and repel growing axons and dendrites (Goodman and Shatz 1993, Tessier-Lavigne and Goodman 1996), and how molecular concentrations can pattern an area to set up pre- and post-synaptic partners (Flanagan and Vanderhaeghen 1998, Benson, Colman et al. 2001, Sanes and Zipursky 2020). However, there still does not exist a comprehensive identification of cell-surface molecules that specifically distinguishes one neuron’s synaptic connectivity from another.

The *Drosophila melanogaster* mechanosensory neural circuit (**Figure 1a**) is an ideal model system to identify molecules involved in pre-specifying synaptic connections. Each mechanosensory neuron has a complex yet invariant synaptic connectivity pattern that forms independent of neural activity (Ghysen 1980, Kays, Cvetkovska et al. 2014). A single mechanosensory neuron innervates a single identifiable bristle on the back of the fly (**Figure 1b**), so each mechanosensory neuron can be uniquely identified from one animal to another. Each neuron can be physically accessed (**Figure 1c**) (or its attendant glia if desired) for single-neuron RNA analysis (Cvetkovska, Hibbert et al. 2013, Dos Santos, Yu et al. 2019), and genetically accessed using different Gal4 drivers (Neufeld, Hibbert et al. 2011, Cvetkovska, Hibbert et al. 2013, Kays, Cvetkovska et al. 2014, Dos Santos, Yu et al. 2019). The highly stereotyped axonal branching pattern of each mechanosensory neuron (Ghysen 1980, Kays, Cvetkovska et al. 2014) indicates a tight genetic regulation of the synaptic connectivity for each neuron, and allows for a quantifiable anatomical readout of connectivity (Chen, Kondo et al. 2006, Neufeld, Hibbert et al. 2011). Additionally, stimulating a mechanosensory bristle elicits a grooming reflex from the animal, providing a behavioural readout of mechanosensory neural circuit function (Cvetkovska, Hibbert et al. 2013, Kays, Cvetkovska et al. 2014, Dos Santos, Yu et al. 2019). Here, we sequence, characterize, and mis-express different cell-surface molecules in identified mechanosensory and chemosensory neurons to begin to generate a comprehensive list of molecules necessary and sufficient to wire up a hard-wired neural circuit. Instead of sequencing broad classes or clusters of neurons as most transcriptomic studies do, we sequence the same identified neuron repeatedly across animals. This allows us to measure the animal-to-animal variability of a single neuron’s transcriptome, which is hypothesized to be as invariant as the synaptic connectivity that it produces. We also test the contrapositive hypothesis that neurons with different synaptic connectivity patterns (i.e., mechanosensory versus chemosensory neurons) have differences in their transcriptomes; these differences, their cell-surface molecules in particular, dictate their synaptic connections. We pair this transcriptome sequencing approach in parallel with a functional RNAi screen of cell-surface receptors that was performed blind to the sequencing. This way the two datasets serve as unbiased tests to identify molecules involved in a neuron’s wiring, with each dataset independently corroborating the other.

**Figure 1.**
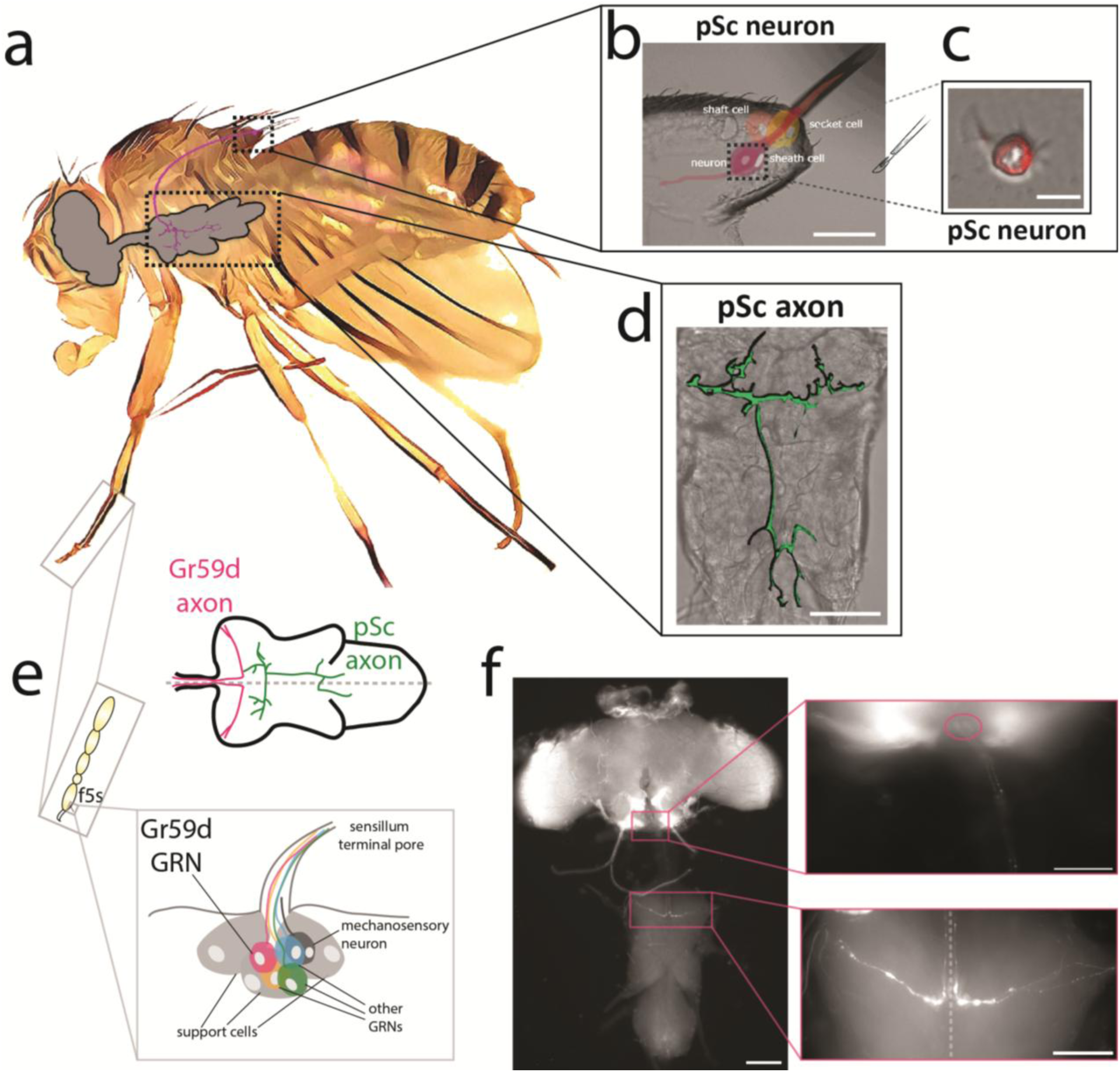
A single, identifiable mechanosensory neuron in *Drosophila melanogaster* has a hard-wired synaptic targeting pattern and can be isolated for RNA sequencing. **a**, The posterior scutellar (pSc) neuron detects movement on the back of the fly and projects into the thoracic ganglion of the central nervous system. **b**, The pSc bristle is composed of the bristle shaft cell, the bristle socket cell, the mechanosensory neuron, the neuron sheath/glial cell. Scale bar is 20μm. **c**, After manual dissection and dissociation, a single pSc neuron expressing a red fluorescent protein was isolated based on fluorescence and morphology. Scale bar is 2μm. **d**, The pSc mechanosensory axonal arbor has a unique and stereotyped wiring pattern within the thoracic ganglion. Scale bar is 50μm. **e,** The Gr59d chemosensory neuron is found on the forelegs of the fly underneath the f5s sensilla. Underneath each sensillum, there are four gustatory receptor neurons (GRNs), one of which expresses Gr59d. There are two f5s sensilla on each foreleg of the fly and therefore two Gr59d chemosensory neurons per leg, and four total in each animal. For clarity, the diagram of the thoracic ganglion shows only one Gr59d axonal projection (magenta), overlaid with the pSc axonal projection (green). **f,** The Gr59d axon does not have as elaborate branching as the pSc arbour. The Gr59d axon enters the thoracic ganglion through the ventral prothoracic neuromere (bottom zoomed image), extends towards the midline but does not cross, and extends up to the brain to terminate in the subesophageal zone (circled in top zoomed image). The scale bar is 100µm for the left panel, and 50µm for the zoomed images. The dotted line is the midline.

## Results

### RNA sequencing of identified mechanosensory neurons

We first sought to compare the molecular differences between the posterior scutellar (pSc) and the anterior postalar (aPa) mechanosensory neurons. Although the pSc and the aPa axons enter the thoracic ganglion within the same nerve fascicle, they each have a distinct synaptic connectivity pattern (**Supplemental Figure 1a**) (Kays, Cvetkovska et al. 2014). Their stereotyped axonal targeting patterns produce important functional differences. Mechanical stimulation of the aPa bristle evokes a grooming behaviour from the animal using its front leg, whereas stimulation of the pSc bristle results in a grooming response from the rear leg (Vandervorst and Ghysen 1980, Kays, Cvetkovska et al. 2014). The exquisite precision, yet similarity, of these neurons’ synaptic targeting patterns argues against molecular gradients as the main determinant of their connectivity differences. Both axons navigate to the same small target region within the thoracic ganglion during pupal development, the period when mechanosensory axons elaborate and refine their arbors (Whitlock and Palka 1995, Venkatasubramanian and Mann 2019, Truman and Riddiford 2023). Across a target field only tens of micrometers wide, a molecular gradient cue would need to change steeply over a few micrometers to precisely specify two different but neighboring arbors. We therefore reasoned that differences in the identity of cell-surface molecules are more likely to distinguish the wiring of these two neurons.

To identify the differences in RNA expression between the pSc and the aPa mechanosensory neurons, we dissected the pSc or aPa bristle sockets in female flies expressing *nSyb-Gal4* and *UAS-mCD8::GFP*, which label all neurons with membrane-bound Green Fluorescent Protein (GFP). We used Fluorescence Activated Cell Sorting (FACS) to isolate and then pool the fluorescent neurons into experimental replicates. We dissected at two time points: the late pupal stage at P14 or adults within a day after eclosion. For each neuron type and each time point, we used a minimum of two experimental replicates containing between ten thousand to thirty thousand sorted cells per replicate. Thus, each replicate represented a single mechanosensory neuron pooled from ∼10,000s of that single neuron (**Supplemental Figure 1b**). RNA was extracted from each of these pooled pSc and pooled aPa replicate samples, and cDNA libraries were sequenced using the Illumina HiSeq 4000 (**Methods**). All neuronal markers (*elav*, *nSyb*, *CadN* and *brp*) were detected in all samples, with an average abundance of 1,278 reads per replicate sample. Glial-specific markers (*Repo*, *moody*, *ana*, and *nazgul*) had very low abundance (< 5 reads per sample). The average number of genes per “neuron” was 10,641 ± 2,674 standard deviation (S.D.). The *Drosophila melanogaster* genome has ∼14,000 protein coding genes. The large number of genes expressed per “neuron” is a consequence of pooling tens of thousands of the same neuron. In other words, low level stochastic expression of a few genes in each neuron (for example, 3 genes) would result in 3,000 extra random genes in 1,000 pooled samples, and a single erroneously transcribed gene in each pSc neuron would result in 10,000 random genes in 10,000 pooled pSc samples. Each mechanosensory bristle has a single sheath/glial cell ensheathing (i.e., physically attached to) the single mechanosensory neuron. The low number of < 5 reads of glial-specific markers per 10,000 neurons strongly rule against the possibility of contamination as a source of variability in gene expression. This is the first time to our knowledge that the same identifiable single cell has been repeatedly sequenced.

Hierarchical clustering revealed significant differences in the transcriptome profiles of the aPa pupal and adult and pSc pupal and adult neurons (**Supplemental Figure 1c**, **Supplemental Table 1**). When the four pooled samples (pSc pupa, pSc adult, aPa pupa, and aPa adult) were clustered, they grouped by developmental age rather than by neuron type: the two pupal samples were most similar to each other, as were the two adult samples. Pupal neurons also showed greater gene expression diversity than adult neurons (**Supplemental Figure 1b**). In aPa neurons, gene expression decreased from 3,300 to 1,500 aligned reads per gene in pupa versus adult, respectively. The pSc neuron decreased its gene expression from 3,800 aligned reads per gene in pupa to 1,900 in adults. In pSc neurons, we identified 10 – 40 genes that were significantly differentially expressed (*p* < 0.05, *n* = 4, *Likelihood Ratio Test*), depending on the type and stringencies of comparisons (**Supplemental Table 2**). We defined differentially expressed genes conservatively as those with an absolute log_2_ foldchange of at least 2 (i.e., a four-fold change) and a Benjamini and Hochberg-adjusted *p* value below 0.05 (**Methods**). All of these genes were more highly expressed in the adult pSc neuron than at the pupal stage. Our results are consistent with previous studies in *Drosophila* neurons that found similar numbers of differential gene expression over developmental time (Li, Horns et al. 2017, Kurmangaliyev, Yoo et al. 2019, Xu, Theisen et al. 2019, Kurmangaliyev, Yoo et al. 2020, Scheffer, Xu et al. 2020, Ozel, Simon et al. 2021, Jain, Lin et al. 2022, Yoo, Dombrovski et al. 2023). These studies and our results show that pupal development is a highly dynamic period for neural transcription, but this two-fold increase in transcription increased the variability in all pupal samples.

Our RNA sequencing data identified 392 total cell-surface molecules and 284 transcription factors expressed in pSc and aPa neurons. Only seven cell-surface or secreted molecule genes were differentially expressed (*p* < 0.05, *n* = 8, *Likelihood Ratio Test*) between pSc and aPa neurons with greater than 10-fold change (**Supplemental Figure 1d, e**, **Supplemental Table 3**), *Eph*, *fat-spondin*, *Tsp42Ed*, *Lar*, *TepII*, *Appl*, and *Tequila*. Five transcription factors were also differentially expressed between pSc and aPa neurons, *Hsf*, *Zif*, *Brk*, *Vsx1*, and *Lz* (**Supplemental Figure 2**, **Supplemental Table 4**). In the *Drosophila* visual system, a combinatorial code of only ten transcription factors can control the identity of nearly 200 neurons (Ozel, Gibbs et al. 2022). We found that using the FACS approach for single neuron isolation yielded a very high quantity of single neurons but produced highly variable RNA data. For example, at the pupal stage, if a single mechanosensory neuron expresses 2–3 incorrectly transcribed genes, then pooling 10,000 of the same neuron would produce 20,000–30,000 extra genes, which is more genes than the *Drosophila* genome. To identify differentially expressed genes involved in neural wiring between different sensory neurons, we used manual dissection and isolation of single sensory neurons guided by fluorescence microscopy for RNA sequencing (**Figure 1c**).

### Single cell RNA sequencing of identified chemosensory neurons

Like mechanosensory bristles, the taste bristles on the leg are arranged in stereotyped positions across different animals, and the chemosensory neurons that innervate these taste bristles also have stereotyped wiring patterns within the central nervous system (**Figure 1e, f**) (Stocker 1994, Scott, Brady et al. 2001, Scott 2018). Only four neurons in the fly express the gustatory receptor Gr59d, and the *Gr59d-Gal4* fly line labels only these four cells (Ling, Dahanukar et al. 2014). One Gr59d neuron is located in the f5s sensillum, and there are two f5s sensilla on the left and the right forelegs (**Figure 1e**) (Ling, Dahanukar et al. 2014). We isolated single Gr59d chemosensory and pSc mechanosensory neurons for single cell RNA sequencing using *Gr59d-Gal4*; *UAS-mCD8::GFP* or *nSyb-Gal4*; *UAS-mCD8::GFP* adult female flies, respectively, to guide the manual dissections of the fluorescent neurons.

Six Gr59d chemosensory neurons and five pSc mechanosensory neurons passed quality control checks for RNA, cDNA, and sequencing. All samples expressed the neuronal gene *elav* and had less than 20 raw counts for the glial genes *Repo* and *moody*. All neurons came from day-old adult female flies, and we detected less than 2 raw counts per sample of five male-specific genes (*Pp1-Y2*, *Ppr-Y*, *kl-2*, *kl-3*, and *kl-5*) in some of our samples. After we normalized the raw counts for sequencing depth and RNA composition, the glial markers and male-specific genes were no longer present, and the *elav* neuronal marker averaged approximately 300 normalized counts. Principal component analysis (PCA) performed on the 11 neurons showed that pSc neurons had higher variability than Gr59d neurons, which was confirmed in the Pearson’s correlations between all pairwise combinations (**Figure 2a, b, Supplemental Figure 3**).

**Figure 2.**
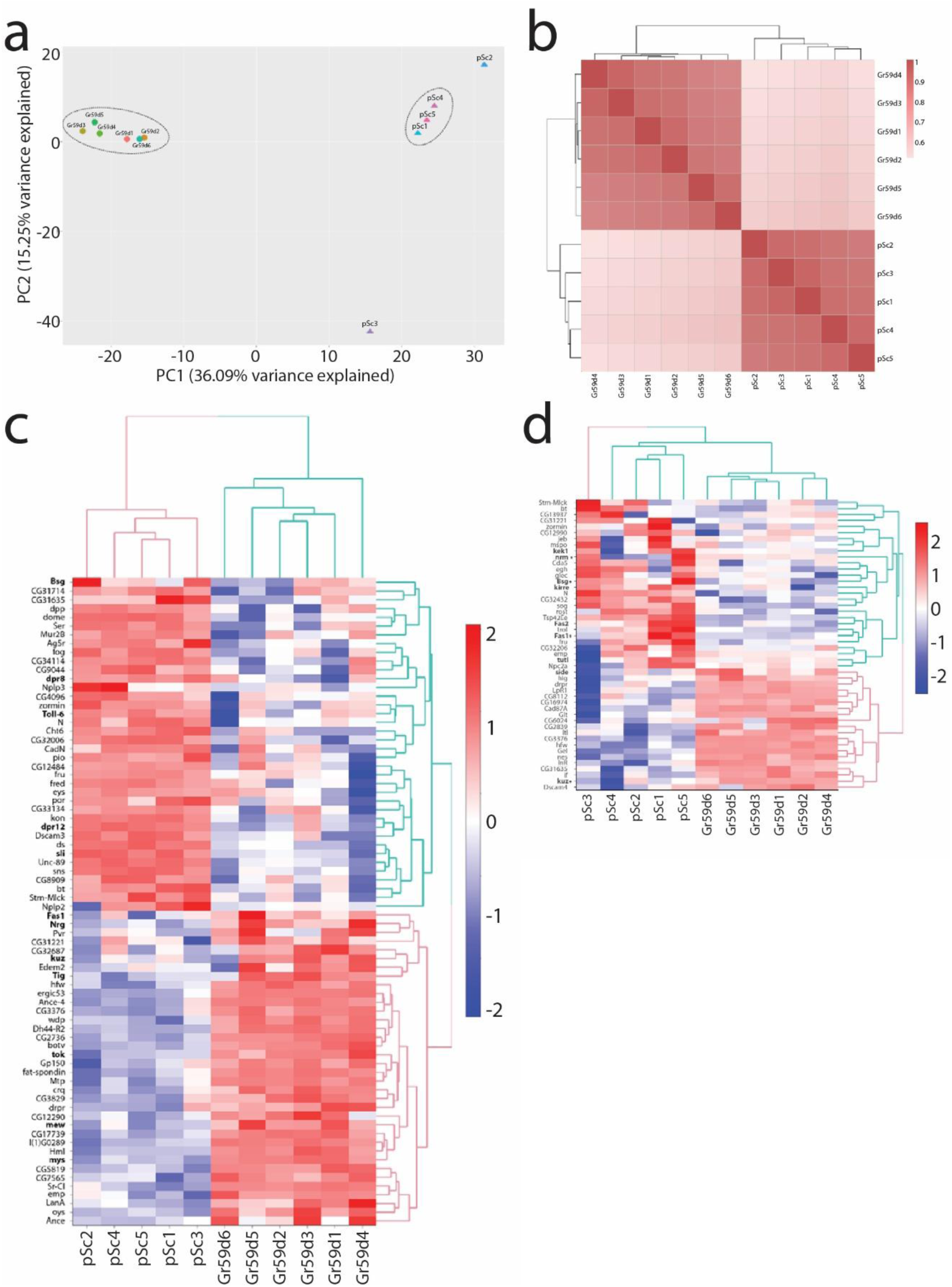
Between 50–70 cell-surface molecules are significantly differentially expressed between pSc and Gr59d neurons. **a**, Principal component analysis (PCA) for adult pSc mechanosensory and adult Gr59d chemosensory neurons. The PCA shows the Gr59d neurons in a tight cluster and away from the pSc neurons. The pSc neurons do not all cluster together and have more variability within themselves. **b,** Pearson’s correlation heatmap for adult pSc and Gr59d neurons also shows a high intragroup correlation for Gr59d neurons and a larger variability for pSc neurons. The expression levels for all genes using normalized log-transformed counts, with low-count genes filtered out, were used. The scale represents the correlation coefficient, *r*. **c,** 74 cell-surface or secreted molecules were significantly differentially expressed between the adult pSc and the adult Gr59d neurons, shown in a hierarchical heatmap, *p_adjusted_* < 0.05, log_2_ (fold change) of |2|. The scale represents the log_2_ (fold change). **d,** 48 cell-surface or secreted molecules were significantly differentially expressed between pupal pSc and adult Gr59d neurons, shown in a hierarchical heatmap, *p_adjusted_* < 0.05, log_2_ (fold change) of |2|. The scale represents the log_2_ (fold change). Genes with an asterisk were also identified in the adult pSc versus adult Gr59d comparison (**c**).

### 74 cell-surface molecules are differentially expressed between the adult pSc neuron and the adult Gr59d neuron

The expression of between 10 to 160 cell-surface molecules can distinguish between different neuron subclasses within the *Drosophila* olfactory (Li, Horns et al. 2017) and visual systems (Tan, Zhang et al. 2015), but these experiments did not examine single identifiable neurons for RNA sequencing. Based on these previous findings, we hypothesized that around 100 differentially expressed cell-surface molecules can determine the wiring differences between the pSc mechanosensory neuron and the Gr59d chemosensory neuron. To identify the genes that are differentially expressed between the pSc mechanosensory neuron and the Gr59d chemosensory neuron, we performed differential expression analysis using DESeq2 (Love, Huber et al. 2014). Out of the 1,448 genes detected in the samples after low-count genes were filtered, we found that 667 genes were significantly differentially expressed between the pSc and the Gr59d neurons. Among these differentially expressed genes, 365 were significantly more expressed in the pSc mechanosensory neuron and 302 were significantly more expressed in the Gr59d chemosensory neuron (**Supplemental Figure 4**). The complete list of differentially expressed genes is in **Supplemental Table 5**. Next, we focused on cell-surface and secreted molecule genes (Kurusu, Cording et al. 2008), and identified 156 cell-surface and secreted proteins in the samples, with 74 genes significantly differentially expressed (**Figure 2c**). 38 were significantly more expressed in the pSc neuron and 36 were significantly more expressed in the Gr59d neuron. The complete list of these differentially expressed genes is in **Supplemental Table 6**.

### 48 cell-surface molecules are differentially expressed between the pupal pSc neuron and the adult Gr59d neuron

Neural wiring within the thoracic ganglion occurs during the pupal development stage before eclosion into the adult fly. We therefore sequenced five pupal pSc neurons at stage p14 (87-90 hours after puparium formation), soon after the mechanosensory neuron differentiates and extends and elaborates its axonal arbor in the thoracic ganglion, and late enough for the soma to be reliably identified and dissected when sensory bristles become mature (Tyler 2000). However, isolating the pupal Gr59d chemosensory neuron proved to be too technically challenging to produce high quality results, but we still sought to examine the molecular differences between the developmental “wiring” stage mechanosensory neuron and the adult chemosensory neuron. PCA on the adult pSc, adult Gr59d, and five pupal pSc samples showed a separation of the Gr59d sample cluster from the pSc neurons. The pupal pSc neurons had a much higher variability in their transcriptomes (**Supplemental Figure 5**). This may be due to differences in developmental times of each pupal pSc neuron dissected, where a one hour difference between neurons during this period of rapid development can contribute substantially to the variance in transcriptomes. After filtering out low-count genes, we performed differential expression analysis among the six adult Gr59d chemosensory neurons and the five pupal pSc mechanosensory neurons. 315 genes were differentially expressed between the pupal pSc and the adult Gr59d neuron, with 231 significantly more expressed in the pupal pSc mechanosensory neuron and 84 significantly more expressed in the adult Gr59d neuron. The complete list of differentially expressed genes is in **Supplemental Table 7**.

Of the cell-surface or secreted molecule genes, we identified 48 that were significantly differentially expressed; 28 were more expressed in the pupal mechanosensory neuron and 20 were significantly more expressed in the adult chemosensory neurons (**Figure 2d**). These molecular differences may be due to differences in developmental stage or in neuron identity. Given this caveat, we place more biological significance on those molecules that were also identified in the adult pSc versus Gr59d comparison (asterisks in **Figure 2d**). For example, the *kuzbanian* (*kuz*) gene was significantly more expressed in the adult Gr59d neuron than in both the pupal and adult pSc neuron stages. Similarly, the *basigin* (*Bsg*) gene was significantly more expressed in the adult and pupal pSc mechanosensory neuron than the adult Gr59d neuron. Curiously, the *Fas1* gene was more highly expressed in the pupal pSc neuron than in the adult Gr59d neuron, but then decreases so that its expression is significantly higher in the adult Gr59d than in the adult pSc state. The complete list of differentially expressed cell-surface and secreted genes is in **Supplemental Table 8**.

### Single cell sequencing of the same neuron reveals transcriptome precision

How precise is gene transcription? The stochastic nature of transcription generates variation even among identical cell clones (Spencer, Gaudet et al. 2009, Tay, Hughey et al. 2010, Flatz, Roychoudhuri et al. 2011, Li and Xie 2011, Lo, Kays et al. 2015, Kays and Chen 2019). Neurons that have a predetermined identity must be defined by a precise molecular code, particularly for neurons that have invariant wiring patterns such as the pSc neuron (**Supplemental Figure 6**). The pSc neuron produces nearly identical synaptic connections across different animals, which requires an impressive level of precision in gene expression (Ghysen 1980, Chen, Kondo et al. 2006, Hattori, Chen et al. 2009, Neufeld, Hibbert et al. 2011, Cvetkovska, Hibbert et al. 2013, He, Kise et al. 2014, Kays, Cvetkovska et al. 2014, Dascenco, Erfurth et al. 2015, Dos Santos, Yu et al. 2019, Dong, Yang et al. 2023). We used single cell RNA sequencing of the pSc neuron to measure its variability in gene expression. Ten pSc neurons from day-old adult females were sequenced to a depth of approximately 200,000 reads on average per cell. As with all single neuron libraries in this study, each of these ten samples passed the same quality controls; they expressed neuronal markers (*elav* and *nSyb*) and contained negligible (< 20) counts of glial markers (*Repo* and *moody*), making contamination by the physically attached sheath/glial cell an unlikely source of the variability we measured. An adult aPa neuron’s transcriptome identity is shown alongside for comparison (**Figure 3a**, inset). We found that single pSc neurons expressed 4,471 ± 1,943 S.D. genes. 50% of pSc neurons expressed the same 3,600 genes, or in other words had 80% of their transcriptome identical. 70% of pSc neurons had the same 1,500 genes transcribed, or 34% of their transcriptome.

**Figure 3.**
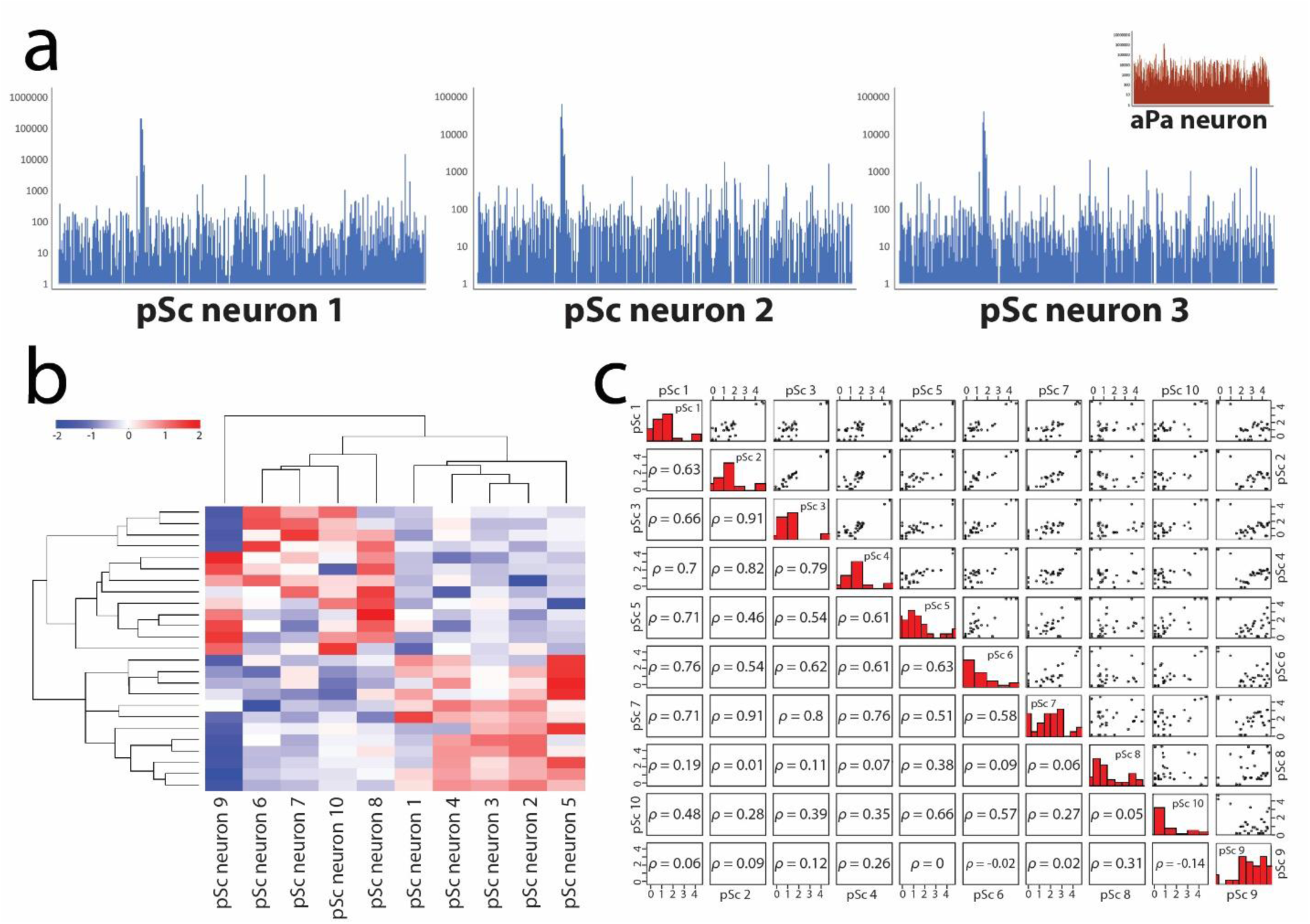
The highly stereotyped connectivity pattern of the pSc mechanosensory neuron is a result of a precise transcriptome. **a**, Three pSc neuron transcriptomes have similar Manhattan Plots. Total reads per gene are shown on the y-axis and genes on the x-axis, arranged alphabetically. The adult aPa transcriptome is shown in the inset for comparison. **b**, Hierarchical heatmap shows few differences in the expression profile of ten single pSc mechanosensory neuron transcriptomes. The scale represents the log_2_ (fold change). **c**, Each mechanosensory neuron has low variability among transcriptomes when compared to each other. The All2All plot displays the correlations and variances between each pSc mechanosensory neuron. The upper right side of the plot shows each log_10_ normalized scatter plot comparing one pSc to another pSc neuron. The bottom left side of the plot is the Pearson correlation coefficient of each pSc pairing. The diagonal shows the histogram of log_10_ normalized read counts. Neurons 8 and 9 were each the least similar to the other eight neurons.

Approximately 1,000 genes were consistently detected in all pSc neurons (**Figure 3a**), and these were also identified in the pooled pSc adult neuron transcriptome. Thirteen genes, or 0.3% of the pSc transcriptome, were significantly differentially expressed (*p* < 0.05, *n* = 10, *Likelihood Ratio Test*) amongst single pSc neurons (**Figure 3b**). Among these thirteen genes, we detected one tumor suppressor (*brat*), one PAX transcription activation domain interacting protein (*Ptip*), one cell-surface receptor (*kek1*), and one transcription factor (*Ef1alpha48D*) (**Supplemental Table 9**). An All2All correlation plot showing the similarities and variances of the ten pSc neurons revealed that pSc neuron samples 8 and 9 were each the least similar to the other eight samples (**Figure 3c**). When comparing pSc neuron 8 to the other nine samples, two mitochondrial genes were differentially expressed, and when comparing pSc neuron 9 to the others, four mitochondrial genes were differentially expressed (*p* < 0.05, *n* = 10, *Likelihood Ratio Test*), indicating that these two neurons were subjected to different environmental factors than the other eight pSc neurons. Transcriptomic variability reveals the level of precision in gene expression that is acceptable to still produce nearly identical hard-wired synaptic connections between animals. Repeated RNA sequencing of an identifiable neuron seeks to address the fundamental nature of variability in connectomics, axonal branching, and cellular identity.

### Single neuron sequencing of alternatively spliced *Dscam1* transcripts

Neuronal identity through a unique transcriptome can also be produced by alternative splicing. Alternative splicing of genes allows for molecular diversity through combinatorial expression of alternate exons of a gene. An extreme example of alternative splicing is the *Drosophila Down syndrome cell adhesion molecule* 1 (*Dscam1*) gene. The *Dscam1* gene can produce tens of thousands of different isoforms through mutually exclusive splicing (**Figure 4a**). It has 20 constitutive exons and 4 variable exon clusters: 4, 6, 9, and 17, where within each of these variable clusters a single exon alternate is spliced to create the single mRNA isoform (Schmucker, Clemens et al. 2000). In *Drosophila melanogaster*, the *Dscam* gene family consists of four independent *Dscam* genes (Millard, Flanagan et al. 2007). The hypervariable alternative splicing only occurs in *Dscam1*, so for simplicity we will refer to *Dscam1* as *Dscam*. The Dscam protein is a cell-surface receptor that is involved in multiple aspects of neural development, neural plasticity, and immune recognition (Schmucker and Chen 2009). Dscam is a large protein and a member of the immunoglobulin (Ig) superfamily, containing ten Ig domains, six fibronectin domains, a single transmembrane region, and a cytoplasmic domain (**Figure 4a**). A single cell is estimated to express between 10 – 50 Dscam isoforms, but these studies were not able to sequence the same identifiable cell repeatedly, and had to sequence each variable exon 4, 6, and 9 independently due to the large size of the *Dscam* mRNA (Hummel, Vasconcelos et al. 2003, Neves, Zucker et al. 2004, Zhan, Clemens et al. 2004). Previous RNA sequence analysis of each variable exon has confirmed that all possible exon alternates are expressed within *Drosophila*, except for pseudo-exon 6.11 (Schmucker, Clemens et al. 2000, Hummel, Vasconcelos et al. 2003, Neves, Zucker et al. 2004, Zhan, Clemens et al. 2004,

**Figure 4.**
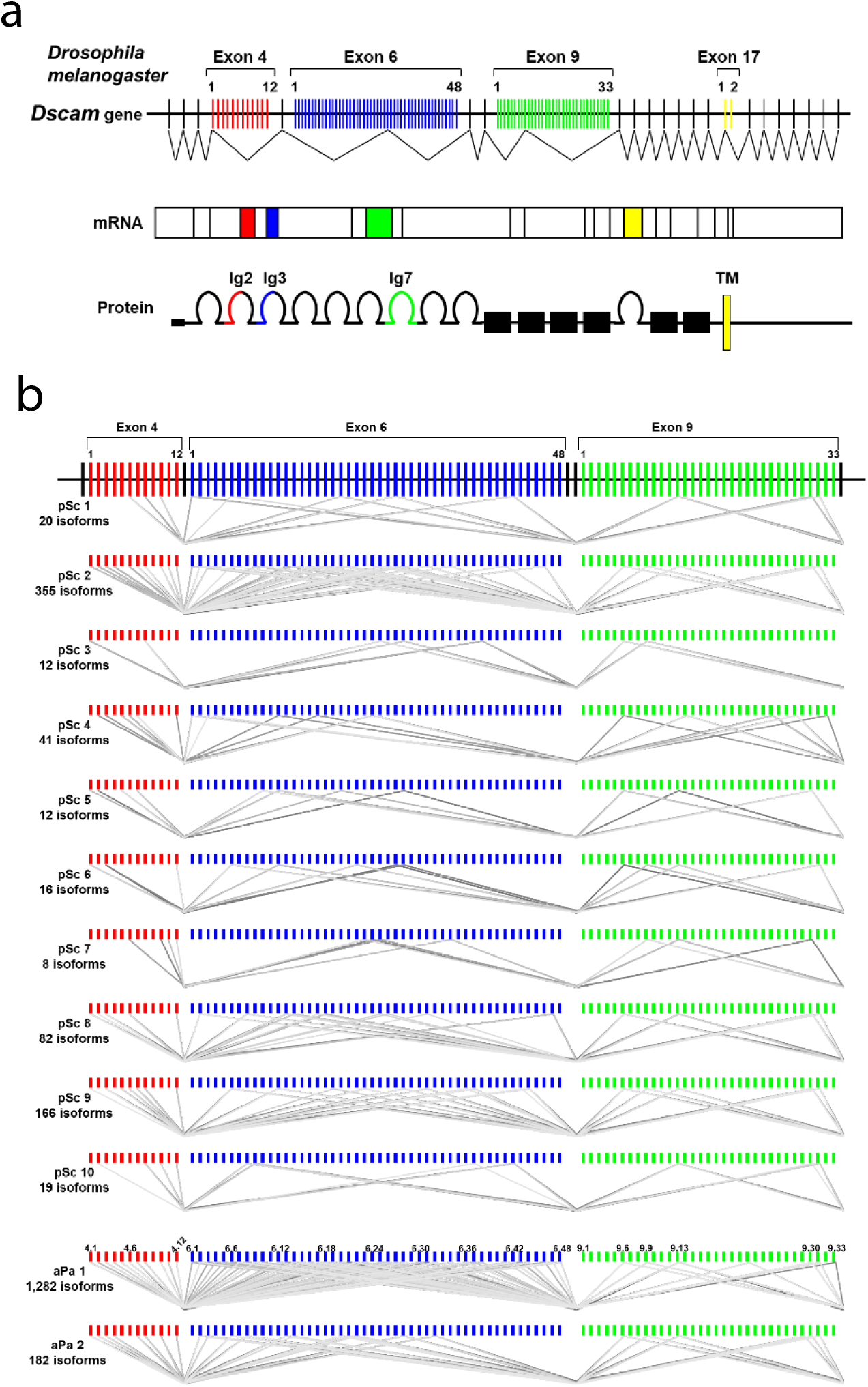
*Dscam* alternative splicing in the pSc neuron is both random and specific. **a**, The *Dscam* gene in *Drosophila melanogaster* exhibits extraordinary alternative splicing in variable exons 4, 6, and 9. Mutually exclusive splicing of exon alternates in these variable exons allows for different mRNA and protein isoforms that maintain the same overall architecture. **b,** Single cell RNA sequencing of ten pSc and two aPa mechanosensory neurons shows preferential splicing for specific exon 9 alternates, 9.6, 9.9., 9.13, 9.30, and 9.31. The aPa mechanosensory neuron had a much higher isoform diversity than the pSc neuron. Grey lines represent individual isoform splicing choices for each neuron; thickness of the line represents the number of reads in the sequencing. Splicing of constitutive exons 5, 7, and 8 are not shown for clarity. Samples pSc1 and aPa1 came from the same animal, and pSc2 and aPa2 came from the same animal.

Schmucker and Chen 2009, Sun, You et al. 2013). *Dscam* splicing has been shown to be regulated temporally and spatially in different tissues, with variable exons 6 and 9 being the most highly regulated (Hummel, Vasconcelos et al. 2003, Zhan, Clemens et al. 2004, Schmucker and Chen 2009, Dong, Yang et al. 2023). For example, a CRISPR screen systematically deleting sections within each variable exon 4, 6, and 9, showed that specific alternates of exons 6 and 9 were required for proper neuronal wiring (Dong, Yang et al. 2023). However, no study has sequenced *Dscam* isoform expression in the same cell between individual animals.

Here, we used PacBio long-read single molecule real-time sequencing on single pSc and aPa adult neurons to determine the reliability of *Dscam* isoform expression within the same cell. We sequenced *Dscam* mRNA isoforms from exon 3 to exon 10 in ten pSc neurons and two aPa neurons from eight day-old adult females with an average number of circular consensus sequencing reads of 26,630 per sample. The pSc neuron expressed on average 73 ± 111 S.D. unique *Dscam* isoforms per cell, ranging from 8 – 355 isoforms (**Figure 4b**). However, the two aPa neurons had a much greater diversity of *Dscam* isoforms, similar to its overall transcriptome, expressing 182 and 1,282 isoforms per cell. The complete list of all *Dscam* isoforms sequenced is in **Supplemental Table 10**.

How precise is alternative slicing? If *Dscam* alternative splicing is deterministic in the same identifiable neuron, this has important implications for its role in neuronal wiring. We found that splicing of variable exon 4 was mostly stochastic (**Figure 4b**), but neither pSc nor aPa neurons expressed an isoform containing exon alternate 4.9. Variable exon 6 exhibited clear biases for specific alternates in both pSc and aPa neurons (**Figure 4b**). For example, seven of the exon 6 alternates were never chosen at all in either neuron. Conversely, an average of 8% and 9% of the isoforms in pSc and aPa neurons, respectively, contained exon 6.28, which should only occur in 2% of isoforms if spliced at random. The average pSc neuron only spliced 10 out of 47 alternates for exon 6, or 21% of the possible choices. The aPa neuron exhibited a more diverse exon 6 splicing profile but had several noticeable differences in the splicing of specific exon 6 alternates compared to the pSc neuron.

We also found a striking amount of precision in alternative splicing in variable exon 9 (**Figure 4b**). Five alternates from exon 9 were spliced consistently among the pSc and aPa neurons, exons 9.6, 9.9, 9.13, 9.30, and 9.31, each with >10% average occurrence among all isoforms. Exon alternate 9.13 occurred on average in 23% and 24% of total isoforms in the pSc and aPa neurons, respectively, making it the most commonly spliced exon 9 alternate. If it were randomly spliced, then the probability of its occurrence should be 1 in 33, or 3%. More than half of all isoforms will contain either exon alternates 9.13 or 9.31. Put another way, all mechanosensory neurons did not splice exons 9.1 – 9.5, 9.7, 9.8, 9.10 – 9.12, 9.15 – 9.21, 9.23, 9.24, 9.26, and 9.29. The probability that a single exon 9 alternate would not be spliced out of 10 isoform splicing events is 74%, out of 50 splicing events is 21%, and out of 100 events is 5%, and out of 200 events is 0.2%. For all mechanosensory neurons sequenced (*n* = 14) out of more than 2,500 isoforms (i.e., unique splicing events) those 21 exon 9 alternates were never spliced. Thus, we found that an identifiable neuron can reliably splice specific *Dscam* isoforms containing specific exon 9 alternates. Variable exon 9 encodes for the entire Ig 7 domain of the Dscam protein, so our results implicate specific properties of Ig 7 variants that may be required for neural wiring or function. Whether any of these exon 6 or 9 splicing specificities are biologically significant remains to be tested functionally. This will require disrupting individual isoforms, for example by using isoform-specific RNAi (Celotto and Graveley 2002) or by CRISPR-Cas9 genome editing to manipulate exon alternates (Dong, Yang et al. 2023).

### The set of cell-surface receptors required to wire up the pSc mechanosensory neuron

To understand how a molecule is used for synaptic targeting, loss of function analyses are required. In parallel with our sequencing experiments, we performed a randomized RNAi screen using the pSc neuron to functionally characterize the cell-surface receptors in the *Drosophila* genome. The masked genes were drawn from a comprehensive literature review (see **Methods**), and not from our differentially expressed gene lists to ensure that the screen was independent of the sequencing results. We sought to identify molecules necessary for creating a mechanosensory neuron’s functional circuit, and keeping the gene identities unknown provided an unbiased verification of the sequencing results. Gene identities were unmasked from the functional RNAi screen and the transcriptomic dataset intersections only after both experiments were complete; therefore, genes identified by both approaches represent the convergence of two mutually unbiased lines of evidence rather than circular validation.

We used the *455-Gal4* driver to express dsRNA solely within the pSc neurons (Kays, Cvetkovska et al. 2014). By imaging the axonal branching patterns within the *Drosophila* central nervous system (**Supplemental Figure 6**), we identified genes involved in different aspects of axon guidance, axonal morphology, and targeting. A total of 214 cell-surface receptor genes were screened (*n* > 10 animals/gene, > 2 different RNAi lines/gene), and 39 of these genes were not detected in our RNA sequencing data (**Figure 5**). Thus, over 80% of the screened genes were expressed in pSc neurons.

**Figure 5.**
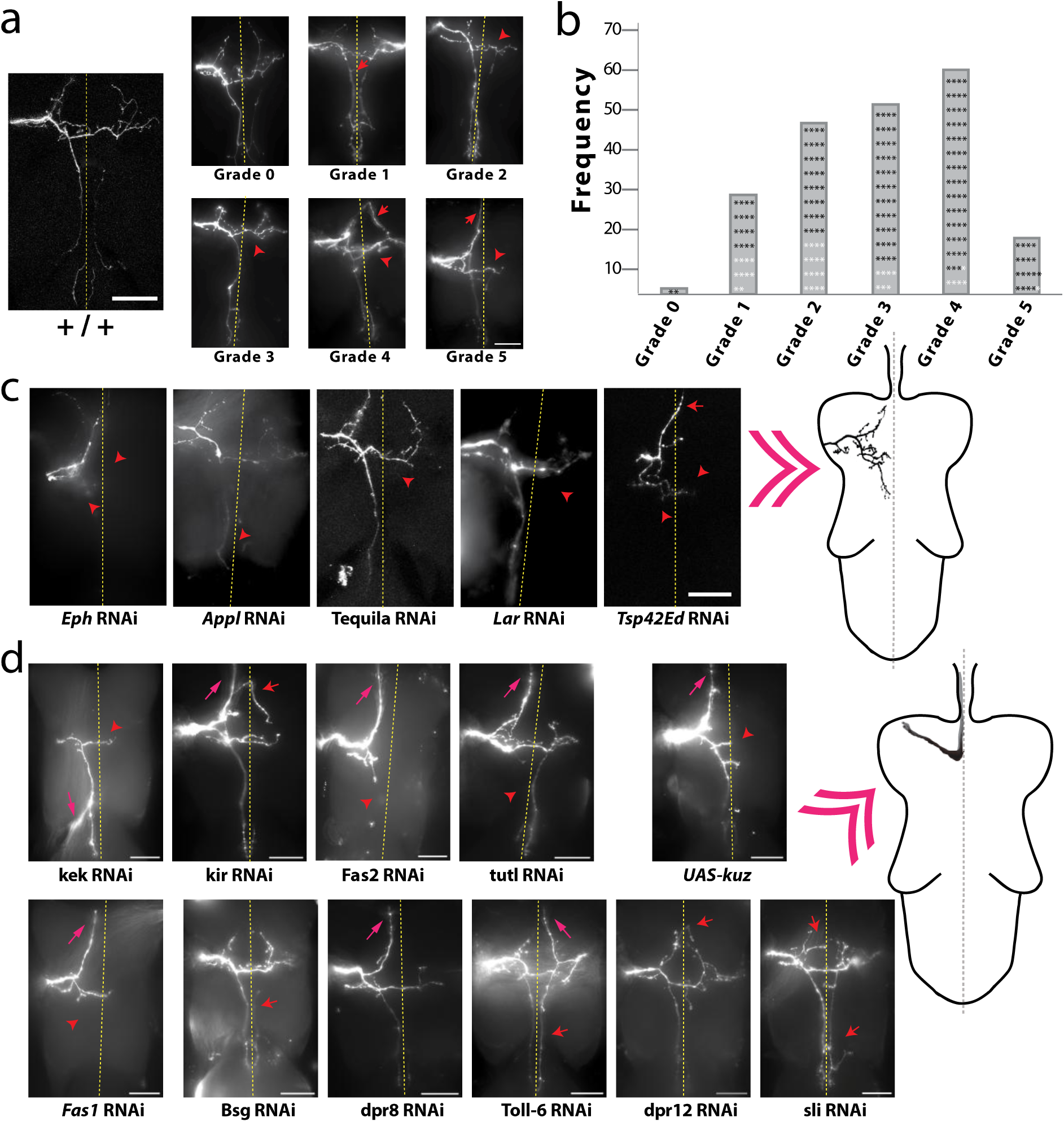
An RNAi screen against cell-surface receptor genes to characterize their specific function in axonal targeting. **a**, Axonal targeting errors were categorized into six grades of severity. RNAi knockdown occurred only within the pSc mechanosensory neuron and experiments and analysis were performed unaware of genotype. Representative examples of pSc axonal arbor phenotypes for each grade are shown. Red arrows point to ectopic branches and red arrowheads indicate missing branches. **b**, Frequency distribution of axonal arbor severity for 214 cell-surface receptor genes, from least severe (Grade 0) to most (Grade 5). Black asterisks indicate a gene that was detected in pSc neurons in the RNA sequencing data; white asterisks indicate genes that were not detected in the RNA sequencing. **c**, Loss of function of differentially expressed genes that were significantly higher in the pSc than in the aPa severely disrupted axonal targeting of the pSc neuron. Representative images of RNAi knockdown mutants of *Eph*, *Appl*, *Tequila*, *Lar*, and *Tsp42Ed* are shown. Red arrows point to ectopic branches, red arrowheads point to missing branches. Schematic on the right depicts the aPa mechanosensory neuron. **d**, Manipulating cell-surface receptor expression within the pSc neuron of differentially expressed genes between the pSc and the Gr59d chemosensory neuron shifted the axonal targeting pattern towards that of the Gr59d neuron (right schematic). RNAi knockdown and mis-expression of differentially expressed cell-surface receptors decreased axonal branch length and complexity and increased axon guidance errors (magenta arrows) of the pSc neuron. Quantification of axonal arbours is shown in **Supplemental Figure 9**. Top row shows RNAi knockdown (left images) and mis-expression (right image, *UAS-kuz*) of differentially expressed genes between the pupal pSc and adult Gr59d. Bottom row shows representative images of RNAi knockdown of differentially expressed genes between the adult pSc and adult Gr59d. The pSc axon exited the thoracic ganglion similar to the Gr59d neuron in nearly all gene manipulations. Red arrows point to ectopic branches, red arrowheads point to missing branches, magenta arrows point to axon guidance errors. Dashed line represents the midline; scale bars are 50μm.

We categorized axonal targeting error phenotypes into six grades. The first three categories included none, subtle, or medium targeting errors (Grades 0 – 2, respectively), and the other three categories included strong to severe axonal errors (Grades 3 – 5) (**Figure 5a**). The average Grade for the 39 genes not expressed in the pSc neuron was 2.3 (**Figure 5b**), which served as randomized negative controls. However, six of the 39 genes, or 15%, that were not detected in pSc neurons in our RNA sequencing (white asterisks in **Figure 5b**) still displayed Grade 4 and 5 axonal targeting error phenotypes. We interpret this 15% as an upper bound on our screen’s false-positive rate for a strong phenotype and reflects RNAi off-target effects and genes that are expressed below the sequencing detection threshold yet remain functional. Based on this background rate, we restricted our functional claims to genes that produced strong to severe axonal error phenotypes (Grade 3 – 5) at high penetrance with at least two independent RNAi lines. Ten genes that we had previously characterized were re-identified in our screen, thus serving as randomized positive controls, and all had strong to severe axonal targeting errors with a penetrance higher than 60%.

Over half of the genes screened (140 genes) showed strong to severe errors with a penetrance higher than 40%. Of these 140 genes, 17 were categorized as Grade 5 and had more than five axonal targeting errors within their primary branch and a penetrance of over 80%. These errors included axonal branch misrouting, ectopic growth, stunted growth, and axon guidance errors (**Supplemental Figure 7**). We further validated our RNA sequencing and RNAi screen using quantitative PCR of pSc neurons. We selected five genes with Grade 5 axonal targeting error phenotypes for single pSc neuron qPCR analysis (*Dpr8*, *Btl*, *Htl*, *Ten-a*, and *Ten-m*), and verified that all five had high levels of expression within the pSc neuron (**Supplemental Figure 8**). Additionally, the *Btl* and *Htl* genes have already been shown to be expressed within pSc neurons using fluorescence *in situ* hybridization (Dos Santos, Yu et al. 2019).

Once the RNAi screen and sequencing experiments were finished, we compared their results and found substantial overlap. For example, five cell-surface molecule genes identified in the screen (*Eph*, *Tsp42Ed*, *Lar*, *Appl* and *Tequila*) were expressed at significantly higher levels in the pSc than in the aPa mechanosensory neuron in our RNA sequencing. RNAi knockdown of these genes resulted in strong to severe axonal targeting errors (Grades 3 – 5) within the pSc neuron (**Figure 5c**). Intriguingly, these error phenotypes reduced the pSc axonal arbor branch lengths and complexities, reminiscent of the aPa axonal arbor. Similarly, nine cell-surface molecule genes that were expressed at significantly higher levels in the pSc mechanosensory neuron than in the Gr59d chemosensory neuron (*Bsg*, *Sli*, *dpr12*, *dpr8*, *Toll-6*, *kek1*, *kirre*, *nrm*, and *tutl*) resulted in strong to severe axonal targeting errors when knocked down (**Figure 5d**, **Supplemental Figure 9**). Loss of function of eight of these nine genes resulted in an axon guidance error where the pSc axon would exit the thoracic ganglion anteriorly to enter the fly brain, similar to the Gr59d axon (**Figure 5d**, **Supplemental Figure 9a**).

### Knockdown of specific cell-surface receptors in the pSc neuron impairs circuit function

Disrupting a sensory neuron’s wiring pattern will disrupt the animal’s perception. We used the pSc bristle cleaning reflex to measure the animal’s response frequency as another assay in our unbiased RNAi screen (**Figure 6a**, **Methods**) (Cvetkovska, Hibbert et al. 2013, Kays, Cvetkovska et al. 2014, Dos Santos, Yu et al. 2019). Selectively reducing each candidate gene’s expression only in the pSc neuron without altering the downstream neural circuitry allowed us to examine how each gene affects pSc function, independent of axonal targeting. We performed the behavioural assay on 49 genes selected from the morphological screen to represent the full range of axonal targeting severities, from no detectable phenotype to Grade 5. This allowed us to test whether, and to what extent, targeting defects lead to a functional grooming deficit (*n* > 10 animals/gene, > 2 different RNAi lines/gene, all perfomed unaware of genotype). A χ^2^ test for independence revealed that only five genes (*Roundabout 1*, *Beat Ib*, *Klingon*, *Beat Ic*, and *Neuromusculin*) were significantly different from control animals (**Figure 6b**). Not surprisingly, the genes whose loss of function resulted in strong to severe axonal targeting errors (Grade 3 – 5, > 60% penetrance), also reduced the animal’s behavioural response frequency (**Figure 6b, Supplemental Table 11**). Interestingly, RNAi knockdown of *Beat Ib* in the pSc neuron resulted in subtle axonal targeting errors, but still significantly reduced the animal’s response frequency, indicating that *Beat Ib* may be involved in synapse function with only a minor role in axonal targeting.

**Figure 6.**
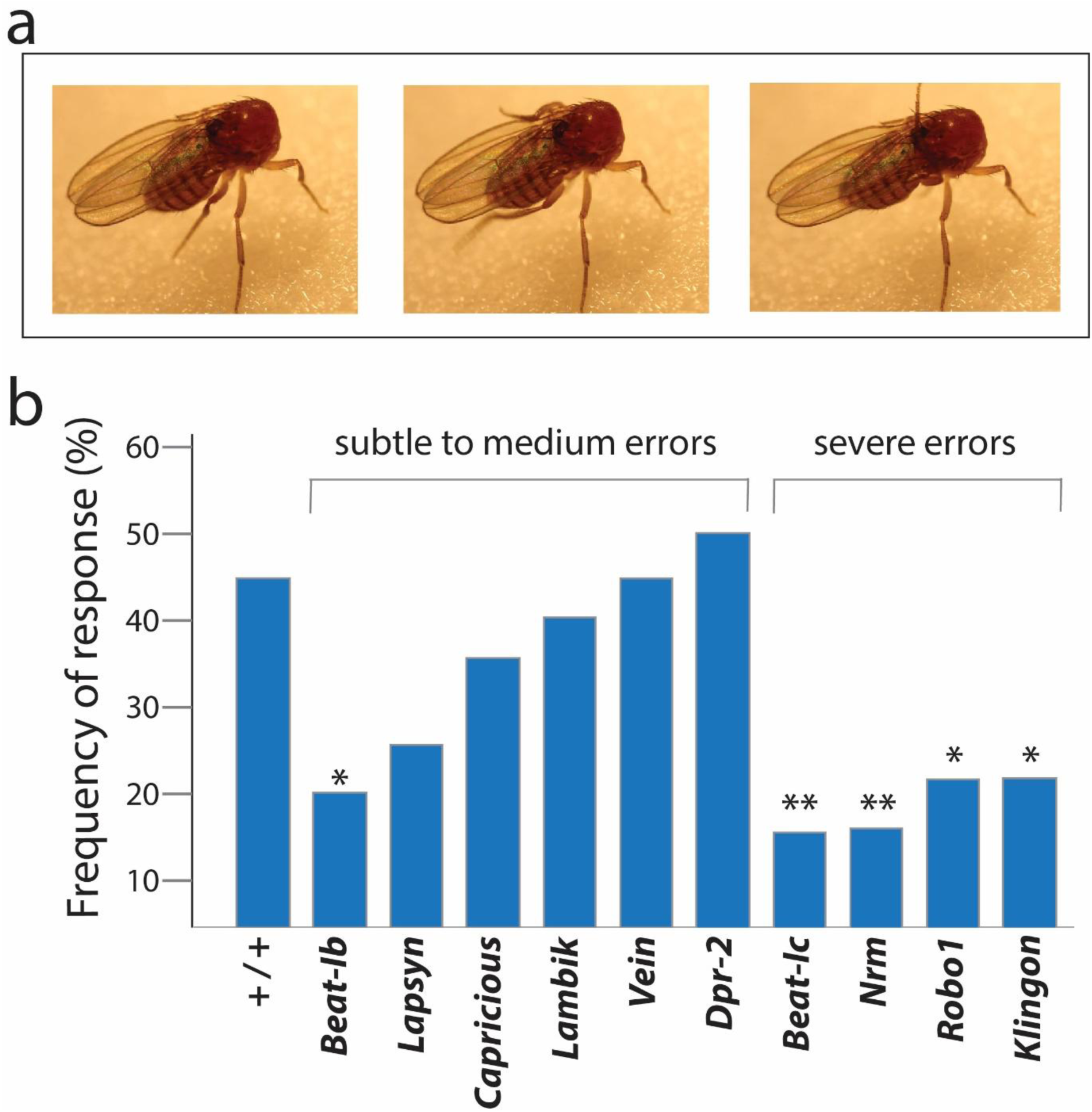
RNAi knockdown within pSc neurons of cell-surface receptors required for synaptic targeting reduces the grooming reflex. **a**, A grooming response from the rear legs of the *Drosophila* can be elicited by stimulating the pSc bristle. **b**, The frequency distribution of animal response rate for grooming versus gene targeted by RNAi is shown. Genes with highly penetrant axonal targeting errors (“severe errors”) in RNAi knockdown (e.g., *Beat-Ic*, *Nrm*, *Robo1*, and *Klingon*) also significantly impaired the grooming reflex when knocked down. Knockdown of *Beat-Ib* selectively in the pSc neuron significantly reduced the grooming response but did not impair axonal arbor morphology. Single asterisk represents *p* < 0.05, double asterisk is *p* < 0.01.

### Re-wiring the Gr59 chemosensory neuron

Loss of function and gain of function experiments can provide evidence for causal relationships between molecules and phenotypes. Based on our RNA sequencing results, we sought to re-wire the Gr59d chemosensory neuron towards the pSc axonal targeting pattern by knocking down and overexpressing specific cell-surface receptors that are differentially expressed. We selected ten genes (*kuz*, *tok*, *Nrg*, *Fas2*, *mew*, *Tig*, *mys*, *Fas1*, *Abl*, and *side*) that were significantly higher expressed within the Gr59d chemosensory neuron compared to the pSc mechanosensory neuron. RNAi knockdown of these genes solely within the Gr59d neuron using *Gr59d-Gal4* resulted in significant increases in ectopic axonal branching within the thoracic ganglion (**Supplemental Figure 10**) (*n* > 9 animals/gene, > 2 different lines/gene).

We also selected eight genes (*dpr12*, *dpr8*, *kek1*, *kirre*, *tutl*, *Toll-6*, *Bsg*, and *Sli*) that were significantly lower expressed within the Gr59d chemosensory neuron compared to the pSc mechanosensory neuron. However, to substantially re-wire a neural circuit, simultaneous knockdown or overexpression of multiple cell-surface molecules is required. Thus, we developed a system to express multiple transgenes simultaneously, under UAS control for conditional expression. We based this system on the mutually exclusive alternative splicing of *Dscam* (**Figure 7a**).

**Figure 7.**
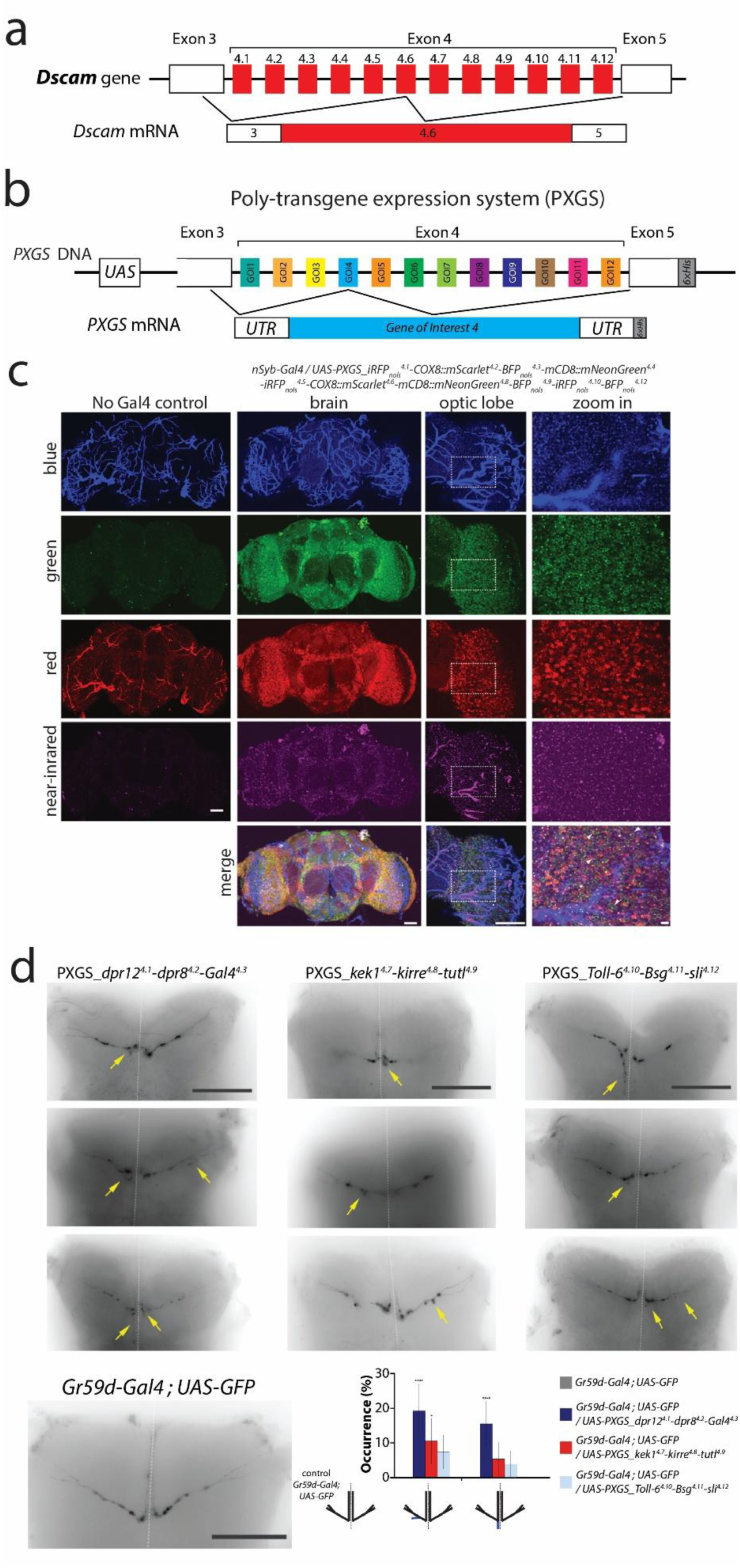
Rewiring the Gr59d neuron. **a**, The mutually exclusive alternative splicing of the *Drosophila Dscam* gene is the basis for the poly-transgene expression system (PXGS). *Dscam* variable exon 4 is shown. **b,** The PXGS system allows for expression of 12 different transgenes by replacing each *Dscam* exon 4 alternate with a gene of interest (GOI). Conditional expression is achieved by placing a *UAS* regulatory sequence upstream of the *PXGS*. Only the last 300 bases of exon 3 and the first 40 bases of exon 5 are kept from the *Dscam* gene, and remain as untranslated regions (UTRs) in the mRNA. A *6×His* tag is included at the 3’ end for ease of molecular biology verification and is also not translated into protein. **c,** After multiple rounds of *PXGS* transcription due to overexpression, *Drosophila* cells express all PXGS transgenes. Transgenic flies containing *UAS-PXGS* constructs with different genes in different alternate positions were used to verify that all possible alternates were expressed. The brains of transgenic flies containing *UAS-PXGS_iRFP_nols4.1_-COX8::mScarlet^4.2^-BFP_nols4.3_-mCD8::mNeonGreen^4.4^-iRFP_nols4.5_-COX8::mScarlet^4.6^-mCD8::mNeonGreen^4.8^-BFP_nols_^4.9^-iRFP_nols_^4.10^-BFP_nols_^4.12^* driven by *nSyb-Gal4* are shown. iRFP is a near-infrared fluorescent protein, nols is a nucleolar localization signal, COX8 is a mitochondrial localization domain, mScarlet is a red fluorescent protein, BFP is a blue fluorescent protein, mNeonGreen is a green fluorescent protein, and mCD8 is a membrane localization domain. White arrowheads in the zoomed in merge image (bottom right) point to cells that express all four fluorophores. **d,** Overexpression of cell-surface receptors differentially expressed from the pSc neuron into Gr59d neurons using PXGS resulted in ectopic axonal branch formation (yellow arrows) within the thoracic ganglion. Representative images of overexpression of *dpr12*, *dpr8*, *kek1*, *kirre*, *tutl*, *Toll-6*, *Bsg*, and *sli* within the Gr59d neuron using *UAS-PXGS* are shown. The frequency of occurrence of two error types, ectopic branching before the midline and midline posterior extension is shown in the bottom right. A schematic for each error is shown below the x-axis, where blue branches indicate ectopic branches. *n* > 6 for all genotypes. Statistical significance compared to control is indicated directly above the bar representing each genotype. Single asterisk represents *p* < 0.05, quadruple asterisks is *p* < 0.0001. The dotted white line is the midline. Scale bar is 5µm in the zoomed in images, and 50µm for all other images.

We extracted the genomic *Dscam* DNA locus starting from the last 300 bases of exon 3, through all of variable exon 4, to 30 bases of the beginning of exon 5, and placed this in a *UAS* vector (**Figure 7b**). We replaced each variable exon 4 alternate 4.1 – 4.12 with a fluorescent protein in a series of transgenic flies to verify that each exon 4 alternate could be replaced, spliced, and expressed properly without any extraneous sequences (**Figure 7c, Supplemental Figure 11**). We called this PXGS (poly-transgene expression system) (Yu, Bucio-Méndez et al. 2025). Using PXGS, we generated three fly lines, *UAS-PXGS_dpr12^4.1^-dpr8^4.2^-Gal4^4.3^*, *UAS-PXGS_kek1^4.7^-kirre^4.8^-tutl^4.9^*, and *UAS-PXGS_Toll-6^4.10^-Bsg^4.11^-sli^4.12^*. Crossing each of these fly lines with *Gr59d-Gal4*, we performed RT-PCR on the Gr59d neuron to verify that the RNA (isolated using the PXGS molecular tag, see **Methods**) for each of the cell-surface receptor genes was expressed in each PXGS position (**Supplemental Figure 12**). Mis-expression of these combinations of cell-surface receptors using *Gr59d-Gal4* resulted in increased Gr59d axonal branching within the thoracic ganglion similar to our RNAi knockdown experiments (**Figure 7d**) (*n* > 10 animals each line). Mis-expression of each of these receptors one at a time within the Gr59d neuron did not significantly alter its axonal targeting, except for overexpression of *kirre* which increased ectopic branch occurrences near the axonal entry point (**Supplemental Figure 10c**). Thus, we demonstrate that knockdown and combinatorial overexpression of specific cell-surface molecules within an identified chemosensory neuron can increase its ectopic axonal branching. Although these manipulations induced pSc-like branching, they did not produce a complete morphological conversion, suggesting that additional differentially expressed molecules may contribute to cell type-specific targeting.

## Discussion

How does the molecular identity of a neuron specify its hard-wired synaptic connectivity? To address this, we combined three approaches in the *Drosophila* mechanosensory and chemosensory systems: repeated RNA sequencing of the same neuron across animals, a functional RNAi screen of cell-surface receptors, and re-wiring of a chemosensory neuron. We found that the transcriptome of an identified neuron varies by less than 1% between animals; that fewer than 10 cell-surface molecules distinguish two mechanosensory neurons with different axonal arbors and that less than 100 cell-surface molecules distinguish a mechanosensory from a chemosensory neuron; that *Dscam* isoform choice in an identified neuron is biased and partly deterministic rather than fully random; that roughly 140 cell-surface receptors are functionally required to build the pSc arbor; and that mis-expressing pSc-enriched cell-surface receptors within a chemosensory neuron is sufficient to induce pSc-like axonal branching. Together these results begin to define a tractable, quantitative molecular code for the hard-wired connectivity of a single neuron, and they provide a single-neuron transcriptome and *Dscam*-isoform resource.

Our experiments took advantage of the unique features of the *Drosophila* mechanosensory system. This system allowed us to identify and isolate the same single neuron at different stages of development. RNA sequencing analysis on the pSc and aPa neurons over time revealed that the differences in transcriptomes were more pronounced between pupa and adult animals rather than between the different neurons and that their transcriptional complexity decreased after the late pupal stage (**Supplemental Figure 1**). These results are consistent with previous single cell analysis studies describing changes in neuronal transcriptomes throughout development (Hanchate, Kondoh et al. 2015, Li, Horns et al. 2017, Kurmangaliyev, Yoo et al. 2020). However, one study found very few significant molecular differences between different clusters of adult *Drosophila* olfactory projection neuron sub-types compared to during the pupal stages (Li, Horns et al. 2017). We identified significant differential gene expression between pSc and aPa neurons even at the adult stage. This difference may be due to the larger axonal arbor size and complexity, or primary sensory function of the mechanosensory neurons compared to olfactory projection neurons or the earlier adult age in our study (1-2 day old versus 3-5 days old). The mechanosensory (and chemosensory) axon extends more than 1 mm from the periphery into the central nervous system and elaborates an arbor across several hundred micrometers synapsing onto hundreds of interneurons. This may require larger concentrations of molecules for synapse function and maintenance far from the soma compared to the slightly smaller second-order olfactory projection neurons within the antennal lobe. Thus, mechanosensory (and chemosensory) neurons may take longer to degrade the higher concentrations of the mRNAs accumulated during synaptic targeting. Or, these neurons may need to stay differentiated and maintain their specific transcriptome and so the cell-surface molecules that we identified also function to maintain synaptic connections, synapse structure, or synapse function.

We found that 99.7% of the pSc transcriptome was not statistically significantly different from each other. However, there are still hundreds to thousands of genes between any two pSc neurons that are different, and if not statistically significant, may be biologically significant. Even if 95% of two transcriptomes were the same, then ∼200 genes are still transcriptionally different. The level of noise in single cell RNA sequencing can come from technical aspects such as ambient RNA, transcript length, RNA priming, library preparation, and sequencing, and biological aspects such as transcriptional bursting, cell stress, cell metabolism, cell state, cell age at time of dissection, and other transcriptional variance (i.e., transcriptional mistakes). Technical aspects can never be ruled out completely as a contribution of variance, but it is unlikely that ambient RNA or contamination is a source of the transcriptional variability that we observed.

The ensheathing glial cell is physically attached to its sensory neuron and we detected very few reads of glial markers per 10,000 samples of single neurons. Our results from RNA sequencing of the exact same identified pSc cell from tens of thousands of isolations of the pSc using FACS and using ten hand-picked pSc samples showed that ∼1,000 genes were consistently expressed in all samples, and the source of cell-to-cell variance was biological (cell stress, cell metabolism, cell state, and transcriptional noise). The remaining variably-expressed genes may not be required for pSc wiring nor function. However, it remains difficult to determine whether outliers in single cell samples from supposedly homogeneous populations are of biological significance (Kolodziejczyk, Kim et al. 2015). The challenges in physically isolating single tiny *Drosophila* neurons from living animals at precise times make it difficult to identify biologically relevant sources of RNA variance. This is more so the case in pupal neurons, where the dissections and isolations are even more challenging and the developmental period is marked by high dynamism and increased transcription and the tissue and cells are more delicate. Our dissections and isolations of tens of thousands of single pSc and aPa mechanosensory neurons took over one year to complete, and minor differences in pupal timing and handling likely contributed to the variance in transcriptomes for the pupal pSc and aPa neurons.

Mechanosensory neurons within the *Drosophila* scutum have axons that all fasciculate within the same bundle and elaborate within micrometers in the same target field of the thoracic ganglion (**Figures 1d, Supplemental Figure 1a**) (Ghysen 1980, Kays, Cvetkovska et al. 2014). We specifically focused on cell-surface receptors and transcription factors that would differentiate between the two neighboring pSc and aPa neurons. Our *a priori* hypothesis did not include intracellular signalling molecules, and transmembrane molecules such as channels, transporters, and pumps. The ∼10 differentially expressed cell-surface molecules and transcription factors are our strongest candidates as determinants of the synaptic connectivity differences between the pSc and aPa neurons. It is not known whether the five transcription factors that differ between the pSc and aPa neurons (*Hsf*, *Zif*, *Brk*, *Vsx1*, and *Lz*) act upstream of the differentially expressed cell-surface molecules. Combinatorial transcription factor codes are known to specify neuronal identity and wiring in *Drosophila* (Ozel, Gibbs et al. 2022), making these transcription factors appealing candidates as regulators of the cell-surface code to test in future work.

Expression data alone cannot establish what a molecule does in a given neuron; each molecule must be functionally characterized in the specific neuron. The 140 cell-surface receptors within the pSc neuron that we identified in our RNAi screen are likely to be generally involved in mechanosensory axonal targeting within the central nervous system, including in the aPa mechanosensory neuron. An aPa-specific Gal4 driver would be required to demonstrate differential molecular functions between the two neurons using genetic manipulations such as mis-expression and loss-of-function. The RNAi screen using the behavioural assay for neural circuit function was far less comprehensive than our pSc neuron morphological screen, due to the large number of animals required for behaviour (∼50 animals per line per gene). However, we identified *Beat Ib*, a cell-surface receptor in the Ig superfamily required for synapse function (i.e., RNAi knockdown of *Beat Ib* in the pSc neuron significantly reduced the grooming response), but not involved in axonal branching or targeting (**Figure 6**). Thus, this combined behavioural and morphological approach can uncover cell-surface receptors such as Beat Ib that may be required for synapse assembly, maturation, maintenance, or active zone recruitment, or vesicle release, independent of synaptic targeting.

One goal of understanding neural circuit formation is to be able to re-wire circuits into designated pathways. We developed the PXGS technique with the aim of expressing 12 different cell-surface receptors and 12 different polycistronic dsRNAs to completely re-wire either the pSc mechanosensory neuron or the Gr59d chemosensory neuron. Our approach to mis-express 3 – 4 cell-surface receptors at a time using PXGS will help determine which combinations of receptors result in specific axonal phenotypes, before expressing all 12 simultaneously. A definitive demonstration of full re-wiring would be behavioural: if a Gr59d chemosensory neuron were functionally re-wired into the pSc mechanosensory circuit, then activating the Gr59d with a bitter tastant should elicit the rear leg grooming reflex normally evoked by the pSc bristle, in essence altering the circuit’s structure, connections, function, and ultimately the animal’s perception. Testing this prediction, for which the PXGS system and our established behavioural assay now provide the necessary tools, is an important future goal.

## Methods

### Fly stocks

To reduce variability due to environmental conditions and genetic background, flies were reared at 25°C on standard cornmeal under 12h light / 12h dark cycles. To express fluorophores in neuronal cells used for RNA sequencing experiments, the pan-neuronal *nSyb-Gal4* driver was crossed to *UAS-mCD8::GFP*, *UAS-tdTomato*, or *UAS-mCherry*. The scutellum specific *455-Gal4* driver line was used to express dsRNA within the pSc mechanosensory neurons (Kays, Cvetkovska et al. 2014). The *Gr59d-Gal4* driver line was used for Gr59d neuron-specific expression (Kwon, Dahanukar et al. 2014). A Gal4 driver for the aPa mechanosensory neuron does not exist. Genotypes denoted as *+ / +* are *455-Gal4 / +*.

A comprehensive review of the literature was done to define cell-surface receptors for the RNA interference (RNAi) screen (Pipes, Lin et al. 2001, Nakamura, Baldwin et al. 2002, Alvarado, Rice et al. 2004, Kurusu, Cording et al. 2008, Sepp, Hong et al. 2008, Naba, Hoersch et al. 2012, McIlroy, Foldi et al. 2013, Ozkan, Carrillo et al. 2013). UAS-TRiP lines were obtained from the Bloomington Drosophila Stock Center. UAS-dsRNA lines were obtained from the Vienna Drosophila RNAi Centre (VDRC), with two lines from each of the VDRC libraries, GD, and KK, when available. The GD collection insertions are P-element based transgenes randomly inserted in the first three chromosomes, whereas the KK collection have two potential genomic landing sites, 40D3 and 30B3, containing phiC31-based transgenes (Dietzl, Chen et al. 2007, Green, Fedele et al. 2014, Vissers, Manning et al. 2016). RNAi off-target effects were minimized by grading targeting error phenotypes only if at least two different lines exhibited the phenotype. Conversely, false negatives may have occurred due to low or non-ideal expression of the dsRNA, or too stringent error phenotyping criteria. The identities of the genes in the RNAi screen remained unknown until the conclusion of the sequencing experiments. A total of 214 cell-surface receptor genes were screened, with an average of 10 flies per gene imaged. **Supplemental Table 11** summarizes the *Drosophila* genes examined.

PXGS plasmids whose expression in *Drosophila S2* cells had been verified by reverse transcription PCR were used to make transgenic animals. Transgenic *Drosophila* fly lines were generated using PhiC31integrase-mediated transgenesis and plasmid injection was performed by BestGene, Inc (Chino Hills, CA).

### Cell preparation for single cell RNA sequencing

The left and right pSc or aPa bristles were dissected from flies expressing either *nSyb-Gal4* or *Gr59d-Gal4* driving a fluorophore. Female flies at the P14 pupal developmental stage or day-old adults were previously screened for fluorescent intensity from the neuron soma underneath the bristle. P14 pupa dissection was chosen as the period soon after mechanosensory neurons differentiate and can be reliably dissected. Variability in mechanosensory neuron age, and thus its transcriptome, can occur when determining the P14 age or during the dissection time moving from one animal to the next over the course of hours to accumulate enough single neurons. For the pSc neuron, the left and right pScs (two pSc neurons per animal) were dissected. Only the left aPa neuron was dissected per animal. Infrequently, the left aPa neuron and the right pSc neuron were dissected from the same individual (**Figure 5**). For the Gr59d neuron, both Gr59d neurons from both the left and the right forelegs were collected (four Gr59d neurons per animal). All dissections were performed at approximately 1PM to reduce time of day or circadian variances in the transcriptomes.

After dissection, 100 μL of 1 mg/mL collagenase in Rinaldini Solution (100 mL H2O, 800 mg/L NaCl, 1200 mg KCl, 800 mg/L NaHCO3, 240 mg/L glucose, 1% Bovine Serum Albumen, pH 7.3) was added to the dissected cells in 200 μL of Supplemented Schneider’s medium (Schneider’s Insect Medium, Fetal Bovine Serum 10%, Streptomycin 2%, and Insulin 0.02 mg/mL) on ice. 100 µL of 10X TrypLE was used to enzymatically dissociate the cells by incubating in a metallic bead bath at 37°C for 15 minutes. The reaction was stopped by adding 500 μL of cold Supplemented Schneider’s medium, and the final cell suspension was mechanically dissociated using a 1000 µL pipette tip and then strained using a 35 μm cell strainer.

For single cell mRNA sequencing experiments, the cell suspension was immediately used for neuronal isolation based on fluorescence intensity and cell morphology using a mouth pipette and an AxioScope A1 epifluorescence microscope (Carl Zeiss) as previously described (Kays and Chen 2019). Single, selected fluorescent cells were pipetted into 5 µL of 1X NEBNext Cell Lysis Buffer from NEBNext® Single Cell/Low Input RNA Library Prep Kit for Illumina on ice before proceeding to cDNA library preparation. For single neuron pooled experiments, pSc and aPa neurons were isolated based on fluorescence intensity using a BD FACSAria™ Fusion sorter (Biosciences) to a total amount of 10,000 to 30,000 events per sample. About 0.1% of dissociated cells expressed the required fluorescent markers and were collected in 200 μL of cell lysis buffer with RNase inhibitor. After isolation, all cells were kept overnight in −20°C.

### cDNA library preparation and sequencing

Full length poly(A)-tailed RNA was used for reverse transcription and amplification using the NEBNext Single Cell / Low Input RNA Library Preparation Kit (New England Biolabs). To increase cDNA yield, we performed 21 cycles of PCR for single cell samples and 14 cycles of PCR for bulk samples. Fragmentation was performed for 25 minutes at 37°C, then 30 minutes at 65°C such that fragments of final library size was between 300-350bp.

Sequencing libraries were prepared using NEBNext Multiplex Illumina Oligos (New England Biolabs). Single cell pSc samples were sequenced using the Illumina HiSeq 4000 PE100 platform with 150bp paired ends, or the Illumina NovaSeq 6000 SP platform with 100bp paired ends. Single Gr59d chemosensory neurons were sequenced using the Illumina NovaSeq 6000 SP platform using 100bp pair-end reads. Pooled pSc or aPa neuron samples were sequenced using the Illumina NovaSeq 6000 SP platform with 100bp paired ends. For *Dscam* isoform RNA sequencing, *Dscam*-specific reverse primers from exon 11 were used for reverse transcription.

We then performed two rounds of nested PCR for 30 cycles each and ligation of SMRTbells for 2.3 kb sequencing. Long read length single molecule real-time (SMRT) sequencing was performed on the PacBio RS platform (Pacific Biosciences). PacBio SMRT sequencing was also verified using 6.6kb direct sequencing where the same *Dscam* isoform combinations and numbers were found. *Drosophila S2* and *Kc* cells were also sequenced in parallel as single cell controls. Intriguingly, the *Kc* cell sample contained only one *Dscam* mRNA isoform that skipped variable exon 9 entirely, resulting in an in-frame mRNA that would produce a Dscam protein lacking the Ig 7 domain (**Supplemental Table 10**). We performed RT-PCR on *Drosophila* tissue, *S2* cells, and *Kc* cells, and verified that this variable exon 9 does occur in cell lines (**Supplemental Figure 13**).

Alternative splicing of *Dscam* variable exon 4 was mostly random, but we found that the pSc and aPa mechanosensory neurons (and a very similar mechanosensory neuron, the aSc, anterior Scutellar neuron, **Supplemental Table 10**) did not splice the exon alternate 4.9.

Because our PXGS technique relies on splicing of *Dscam* variable exon 4, it is important to characterize which exon 4 alternates are expressed in the cells of interest. We verified that the Gr59d neuron expresses all PXGS exon 4 alternates by crossing to different *UAS-PXGS-fluorophore* flies and checking for fluorescence.

### RNA-sequence quantification and analysis

Reads were trimmed from the 3’ end to have a PHRED score of at least 30. Illumina sequencing adapters were removed from the reads, and all reads were required to have a length of at least 32 bp. Trimming and clipping was performed using Trimmomatic (Bolger et al., 2014). Reads were aligned to the *Drosophila melanogaster* genome (BDGP6.22) using STAR (Dobin, Davis et al. 2013) (v2.7.3a) with default settings. Successfully aligned uniquely mapped reads were counted using HTSeq-Counts (Anders, Pyl et al. 2015) (v0.11.2) and organized into a raw read counts table. Alternatively, alignment and counts were also performed in Kallisto (Bray, Pimentel et al. 2016) (v0.46.1) and Seurat (Stuart, Butler et al. 2019) (v3.1.0).

All subsequent analysis (principal component analysis and differential expression analysis) was performed in R (v4.2.1) using edgeR (Robinson, McCarthy et al. 2010) (v3.28.0), limma (v3.42.0), Glimma (Law, Alhamdoosh et al. 2016) (v1.14.0), DeBrowser (Kucukural, Yukselen et al. 2019) (v1.14.1), and DESeq2 (Love, Huber et al. 2014) (v3.15). Hierarchical clustering was performed using the R package pheatmap. All packages are available at bioconductor.org. The raw read counts table was filtered such that mtRNA (37 genes), tRNA (320 genes), srRNA (102 genes), snRNA (104 genes), snmRNA (36 genes), snoRNA (284 genes), scaRNA (6 genes), and sometimes miRNA (236 genes) and mitochondrial genes (37 genes) were removed before analysis. Low-count genes were filtered using the counts-per-million (CPM) method (CPM threshold of 1).

The processed expression matrix was subjected to several preprocessing steps. Genes that were expressed in less than 10 cells were removed. Cells were also filtered based on the number of detected genes, number of detected unique molecular identifiers (UMIs), house-keeping gene expression and percentage of mitochondrial gene expression. Cells that expressed less than 10 house-keeping genes were removed. For UMIs, detected genes and mitochondrial gene expression, cutoffs were defined as the more conservative value between a hard predefined cutoff (UMI: lower 1,000 – upper 20,000; genes: lower 200 – upper 5,000; mitochondrial gene expression: 10 percent) and a dataset specific cutoff computed using interquartile ranges. Surface proteins were normalized using the centered-log ratio (CLR) method.

For differential expression analysis, a parametric fit was used for the dispersion type. The Wald significance test was performed and differentially expressed genes were defined as any gene that had log_2_(foldchange) of |2| with a corrected *p* value of less than 0.05. The *p* values in the Supplemental Tables represent the family-wise significance level, or multiplicity adjusted *p* value (e.g., pSc pupa vs aPa pupa vs pSc adult vs aPa adult). The Benjamini and Hochberg method was used to obtain the corrected *p* value considering the effect of multiple testing.

Median ratio normalization was applied to normalize the count data before generating the heatmap of differentially expressed genes and the boxplots for the individual genes. For differential expression analysis of cell-surface molecules and secreted genes only, the raw read counts table was filtered against a known list of 971 *Drosophila* cell-surface molecules and secreted genes, and only read counts for these cell-surface molecule genes were used for differential analysis as described above (Kurusu, Cording et al. 2008). A curated list of *Drosophila* transcription factors from the FlyTF v1 database was used. For the ten single neuron RNA sequencing libraries, single cells were sequenced to a depth of ∼200,000 reads per cell on average, and ∼1,000 genes were consistently detected in all samples after excluding genes with 0 reads for more than one sample. For the pooled single pSc or aPa neurons, four different categories were used: pSc pupa, pSc adult, aPa pupa, and aPa adult, each in experimental duplicate. The sequencing datasets are located at https://www.ncbi.nlm.nih.gov/bioproject/1219304, and the accession number is: PRJNA1219304.

### Quantitative Polymerase Chain Reaction (qPCR)

The procedure for cell preparation of the left and right pSc neuron was identical for single cell RNA sequencing, described above. Thirty cells were examined in all. 2 μL of single cell lysate was used per 20 μL of qPCR reaction using Cells-to-CT™ 1-Step TaqMan™ kit and custom designed qPCR probes for *Dpr8*, *Breathless*, *Heartless*, *Teneurin-a*, and *Teneurin-m*.

Each experimental replicate compared the expression level of all five genes in the same cell to *Drosophila* genomic DNA (200 ng/μL). Gene expression results were quantified as the ΔCT (cycle threshold) compared to the genomic DNA control. A 96-well plate was loaded in a StepOnePlus™ Real Time System (ThermoFisher) according to the long cycle settings described by the manufacturer.

### Generating Poly-transgene expression system (PXGS) constructs

The entire variable exon 4 region of the *Dscam* gene was amplified from *Drosophila melanogaster* genomic DNA from 300 bp from the 3’ end of exon 3 to 40 bp after the 5’ end of exon 5. This *Dscam* fragment was then inserted into the *pJFRC7-20XUAS-IVS-mCD8::GFP* expression vector (Addgene, Cambridge, MA) by replacing the *mCD8::GFP* sequence. A *6×His* tag was added at the end of exon 5 to distinguish PXGS sequences from endogenous *Dscam* sequences in subsequent experiments. To be clear, each transgene has its own start and stop codon at each exon 4 alternate position, so the exon 3 and exon 5 sequences are transcribed but not translated and act as 5’ and 3’ untranslated regions, and this is the case for the *6×His* tag as well; it is transcribed as an RNA tag but is not translated and will not interfere with protein function.

Poly-transgene constructs were into the *pJFRC7-UAS-PXGS* plasmid. Molecular assembly strategies for transgene insertions into the PXGS exon alternates were generated using the automated Molecular Assembly feature in GeneDig.org (Suciu, Aydin et al. 2015), primarily using gene synthesis, Gibson assembly, and restriction enzyme digest and ligation. Exon 4 alternates were replaced by fluorophores or cell-surface receptors. The *BFP_nols_* gene (blue fluorescent protein with a nucleolar localization signal) was obtained from a plasmid previously generated in our lab (Lo and Chen 2019). The *iRFP_nols_*, *mCD8::mNeonGreen*, and *COX8::mScarlet* genes were codon optimized for *Drosophila* and synthesized by Integrated DNA Technology (Coralville, IA). Sequences for cell-surface receptors were obtained from RT-PCR on wildtype flies. All DNA constructs were verified by colony PCR and Sanger sequencing.

### Carbocyanine Dye Labeling and *Drosophila* imaging sample preparation

*Drosophila* mechanosensory axonal arbor morphology was assessed using carbocyanine dye injection into the open sockets of pSc neurons as previously described (Chen, Kondo et al. 2006, Neufeld, Hibbert et al. 2011, Cvetkovska, Hibbert et al. 2013, Kays, Cvetkovska et al. 2014, Dos Santos, Yu et al. 2019). The left and right pSc bristles of 2-day old female flies were plucked and the thorax was left overnight in fixative solution (3.7% paraformaldehyde in 0.2 M carbonate-bicarbonate buffer, pH 9.5). On the next day, the left and right pSc neurons were labelled using fluorescent carbocyanine tracers, DiI or DiD (ThermoFisher) dissolved in ethanol (at 20 mg / mL and 40 mg / mL, respectively). The dye-filled flies were left undisturbed and protected from light in a Petri dish, partially immersed in 0.2 M carbonate-bicarbonate buffer at pH 9.5. After 48 hours, the thoracic ganglion was dissected and placed ventral side up on a slide and mounted with a coverslip.

For *ex vivo* imaging of the Gr59d chemosensory axon, 2-day old female flies expressing *Gr59d-Gal4* driving *UAS-mCD8::GFP* were used. Animals were anesthetized on a CO2 pad and the thoracic ganglion was dissected and mounted ventral side up for immediate fluorescent imaging.

### Image acquisition and analysis

Fluorescent images were acquired using an AxioScope A1 epifluorescence microscope (Carl Zeiss) with a 40× objective (N.A. 1.0) or an Olympus laser-scanning confocal microscope FV1000 with a 40× objective (N.A. 1.3). Within any set of directly compared axonal arbors, images were acquired on the same microscope with identical acquisition settings. 10–20 images of the axonal arbor were taken at different focal planes. Maximal intensity projections of the images were stacked in the Z dimension to create a final image. Images were only adjusted for contrast and brightness when required. For presentation purposes, pSc axonal arbours are shown as white on a black background, and Gr59d arbours are inverted as black on a white background. Transmitted light images were acquired and used to evaluate dissection and dye labeling quality and to measure thoracic ganglia width. Only images without central nervous system damage or surface occlusions were included for data analysis. Measurements of axonal branch length and branch number were performed unaware of genotype by randomly distributing the axonal arbor data of the total dataset. Custom-written software in MATLAB was used to perform the quantitative image analysis.

To determine the baseline variability in wildtype pSc mechanosensory neurons, the total number of branches and branch lengths per arbor were measured from thirty control *455-Gal4 / +* animals. A skeleton of the control *455-Gal4 / +* axonal arbor was then created (**Supplemental Figure 6**) based on the frequency of occurrence of each branch, which was found to be consistent with our previous measurements (Chen, Kondo et al. 2006, Neufeld, Hibbert et al. 2011, Cvetkovska, Hibbert et al. 2013, Kays, Cvetkovska et al. 2014, Dos Santos, Yu et al. 2019). We defined an ectopic branch as any branch that has less than 30% occurrence in the wildtype. For the Gr59d neuron, 23 *Gr59d-Gal4* / + animals were used to determine baseline variability in the control condition. Representative examples of axonal arbors in figures were randomly chosen from the sets of all axonal arbor images of the genotype.

### Behavioral assay and analysis

The *455-Gal4* driver was used to selectively express *UAS-dsRNA* in the pSc neurons to ensure that the RNA interference did not perturb the downstream postsynaptic neural circuitry. Two-day old female flies were decapitated and placed in a humidified chamber to recover over a period of one hour. After recovery, only flies standing upright were evaluated for preliminary responsiveness by eliciting the cleaning reflex from the forelegs after stimulating the anterior notopleural (aNp) bristle with forceps. Responsive flies were then assayed by pressure injection of fluorescent dye onto the posterior scutellar bristles (**Supplemental Video 1**). The presence or absence of a response was scored visually, with a positive response considered as the successful movement of the rear pair of legs towards the labelled bristles with dye spread to the rear legs.

Further description of this protocol can be found elsewhere (Kays, Cvetkovska et al. 2014). A total of 49 cell-surface receptor genes were screened, with an average of 100 flies per gene assayed.

### Statistical analyses

As inclusion and exclusion criteria for the study, single cell RNA sequencing libraries were included only if they passed the neuronal marker, glial marker, and read depth quality control checks described above, and axonal arbors were included only if the nervous system was undamaged and unobscured; no other data were excluded. For the RNAi screen imaging experiments, an initial pilot study was conducted on ten animals per genotype on 50 genotypes to determine adequate sample size through power analysis. Throughout the full RNAi screen, experimenters were unaware of the genotype during the entire process. Targeting error phenotypes were classified into Grades 0 to 5 after quantitative assessment of three criteria: severity of axonal branching pattern change, measurement of the number of axonal branching errors, and penetrance. An image is considered within a Grade if it meets all three criteria for that Grade. If an image fails to meet one criterion, then it is downgraded to the Grade below. **Supplemental Table 12** summarizes the criteria used to classify phenotypes. Statistical significance for total branch length and the number of branches was determined using one-way ANOVA followed by Dunnett’s multiple comparisons test. Statistical analysis was performed using GraphPad Prism version 8.0.0 for Windows (GraphPad Software, San Diego, California USA) and SPSS (version 25). Statistical significance for frequency of occurrence was determined using a two-tailed t-test for proportions in R (version 4.2.1). *F* test for comparison of standard deviation was performed using MedCalc (v20.218). The datasets, analyses, and materials used in the current study are available from the corresponding author upon request.

## Acknowledgements

The authors thank Ibrahim Kays for assistance with experiments. This work was supported by a grant from the Natural Sciences and Engineering Research Council of Canada (2020-04902, B.E.C.). The funding agency had no role in the design of the study and collection, analysis, and interpretation of data and in writing the manuscript.

## Author Contributions

J.V.S., R.Y.Y., and B.E.C. contributed to the conception and design of the work, the acquisition, analysis, and interpretation of data, and drafted the work and reviewed it critically for important intellectual content. R.S. contributed to the conception and design of the work. A.A.T., P.M.P., D.R., V.C., A.B-M., T-J.L., F.E., H.D., P.B. contributed to the acquisition, analysis, and interpretation of data. All authors gave final approval of the version to be published.

