## Supplemental Information for "Identifying molecular instructions to hard-wire a sensory neuron’s synaptic connectivity"

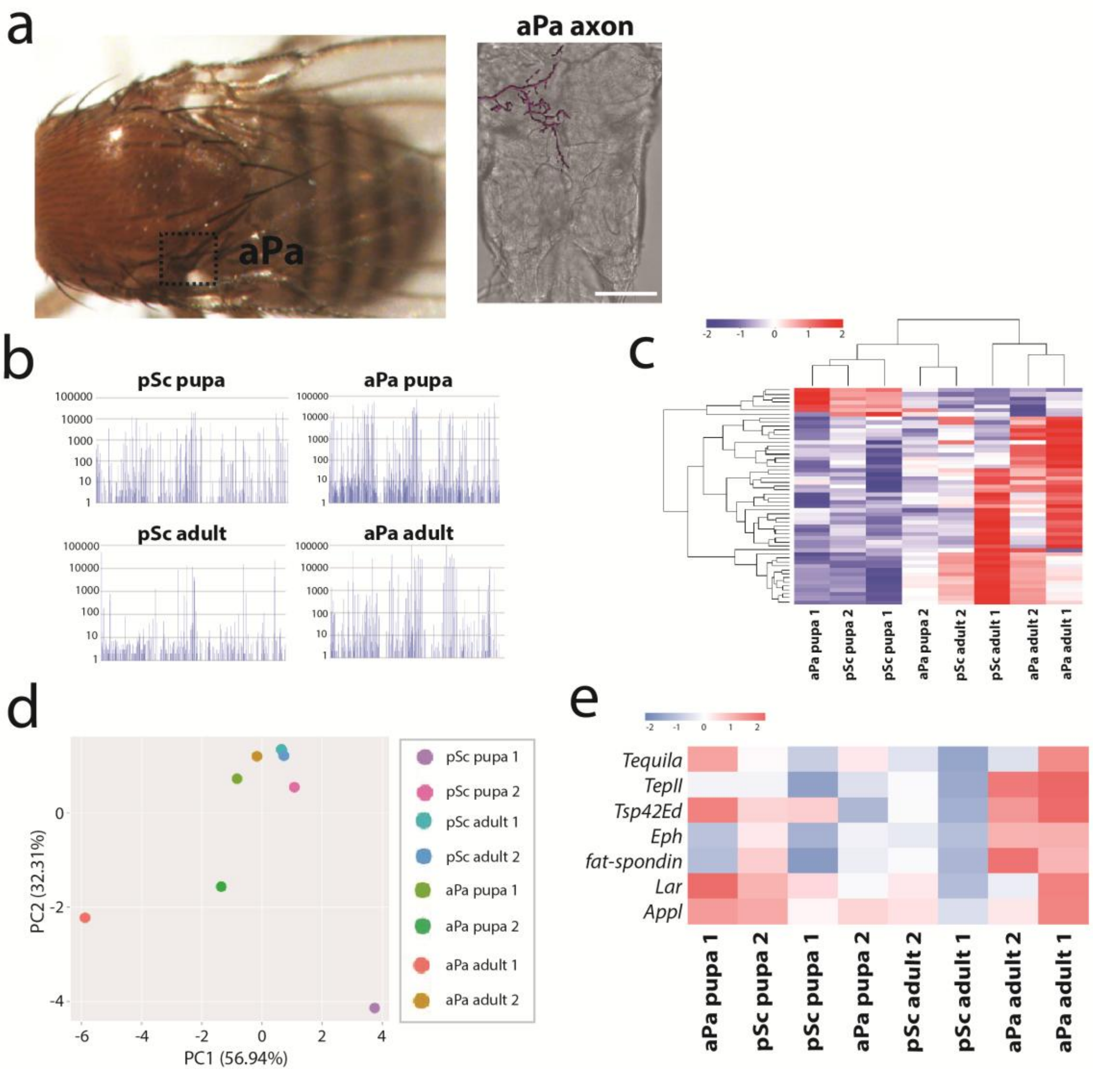


**Supplemental Figure 1 – Approximately ten cell surface receptors can distinguish the pSc mechanosensory neuron from the aPa mechanosensory neuron.** **a,** The anterior postalar (aPa) mechanosensory axon enters the thoracic ganglion within the same fascicle as the pSc axon and has a similar but less complex axonal arbor, and does not cross the midline. The aPa neuron innervates the aPa bristle which neighbors the pSc bristle. Scale bar is 50µm. **b,** Each mechanosensory neuron at each time point has a unique transcriptome. Tens of thousands of dissected pSc or aPa neurons were pooled per replicate and their RNA sequenced to represent the single neuron. A comparative view of total reads (y-axis) per gene (x-axis) is shown in four Manhattan Plots of the pooled samples of pSc pupa, pSc adult, aPa pupa, and aPa adult neurons. The 16,000 genes on the x-axis are arranged alphabetically. **c**, A hierarchical heatmap illustrates the differences in the transcriptional profiles of pSc and aPa mechanosensory neurons at the pupa and adult stages. The scale represents the $\log_2$ (fold change). **d**, PCA of the RNA sequencing data selectively filtered for cell surface receptor genes separates the pSc neurons on the right and aPa neurons on the left. **e**, Seven cell surface receptors are differentially expressed between pSc and aPa mechanosensory neurons, shown in a hierarchical heatmap, $p_{adjusted} < 0.05$, $\log_2$ (fold change) of |2|. The scale represents the $\log_2$ (fold change) between pSc versus aPa neurons.

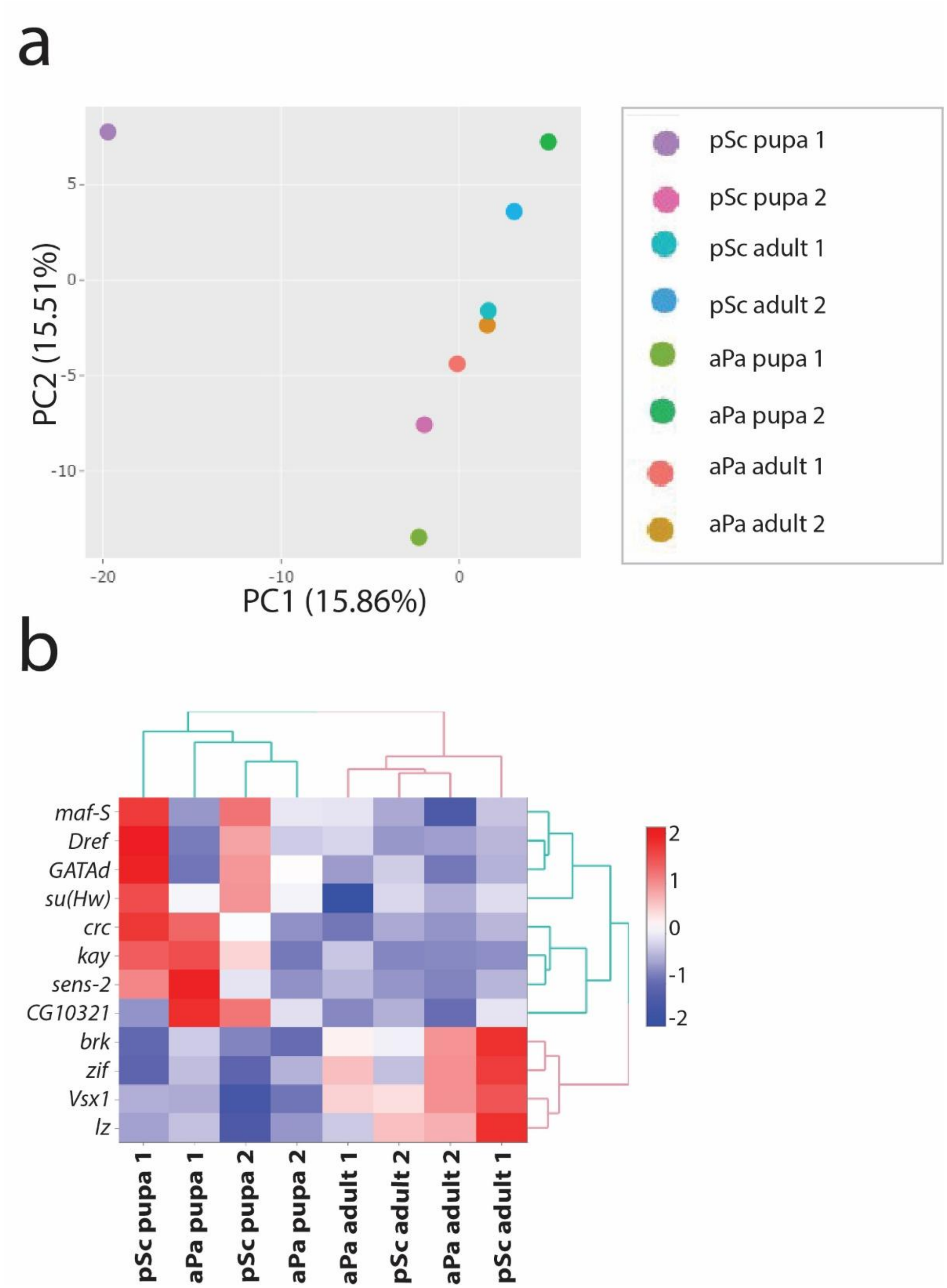


**Supplemental Figure 2 – A small set of transcription factors can distinguish between mechanosensory neurons. a**, Principal component analysis of the RNA sequencing data selectively filtered for transcription factor genes. **b**, Hierarchical heatmap of the differential expression of transcription factors between pSc and aPa mechanosensory neurons at pupa and adult stages, $p_{adjusted} < 0.05$, $\log_2$ (fold change) of $|2|$. The scale represents the $\log_2$ (fold change) between pSc versus aPa neurons.

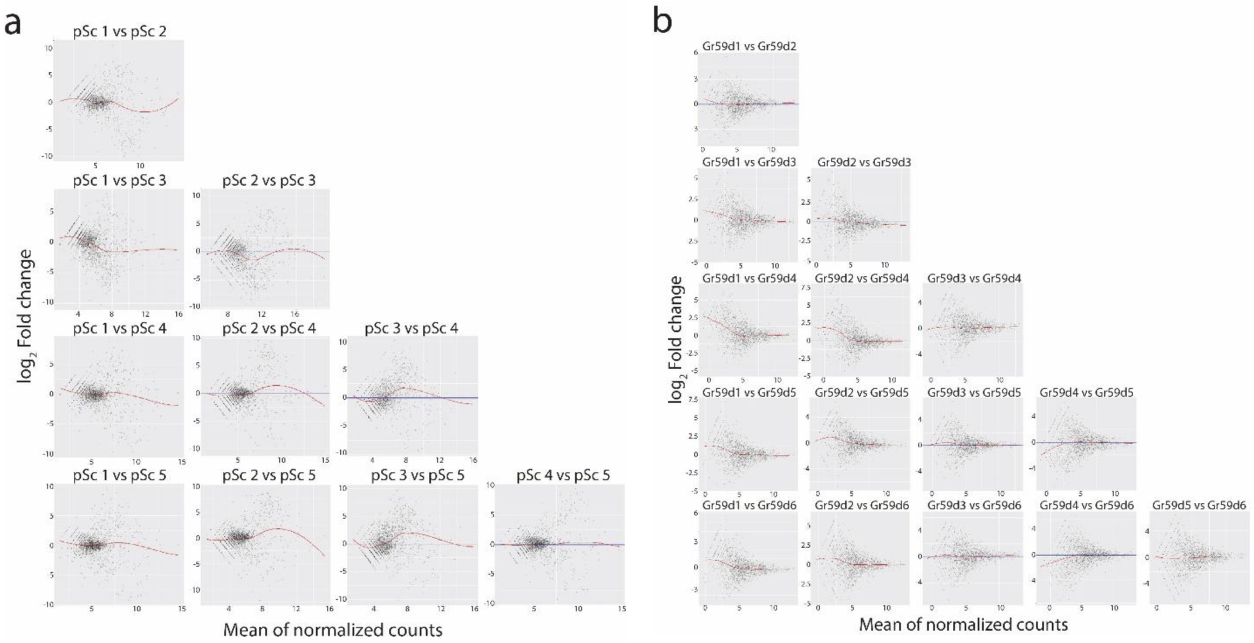


**Supplemental Figure 3 – Gene expression variability between pSc mechanosensory neurons versus Gr59d chemosensory neurons. a, b,** An MA plot compares the log fold change for each gene (**M**, or y-axis) versus the mean of normalized expression count (**A**, or x-axis) for each gene between two samples. MA plots compare the gene expression variability within pSc samples (**a**) or Gr59d samples (**b**). The trend of means was visualized using LOESS (locally weighted smoothing) regression of means, shown in the red regression curves. The more similar the two samples are, the more the data points will converge to zero and result in a flat regression line at M=0. Genes with lower mean expression values (left of each MA-plot) are expected to have more variability in log fold change, thus a not-so-flat regression line (McDermaid et al., 2019). For the pSc samples, except for pSc4 vs pSc5, all other comparisons' regression lines had non-zero slopes, suggesting that the samples being compared have some variance in between. In contrast, for Gr59d samples, most regression lines converged around M=0 as the graph moved right and beyond the low mean expression values. In other words, there is more variance in gene expression between any two pSc neurons than there is between any two Gr59d neurons.

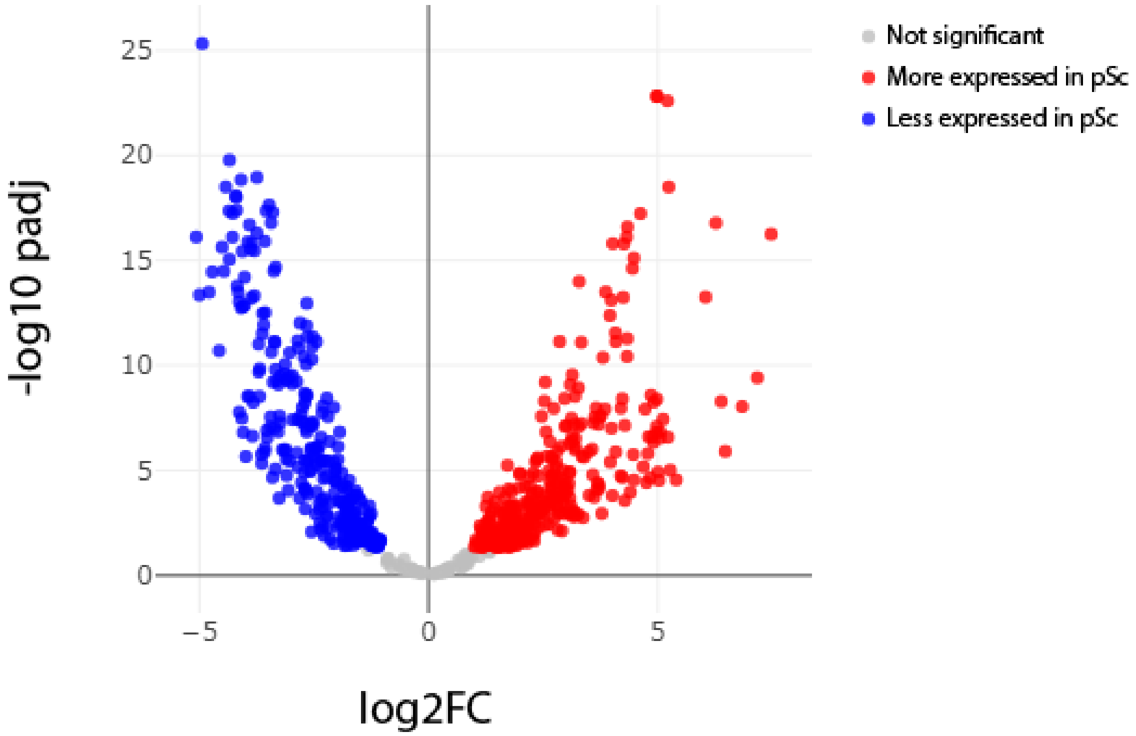


**Supplemental Figure 4 – volcano plot showing that 667 genes were differentially expressed between adult pSc neurons versus adult Gr59d neurons, $p_{adjusted} < 0.05$, $\log_2$ fold change of |2|.** 365 were significantly more expressed in the pSc mechanosensory neuron (red dots) and 302 were significantly more expressed in the Gr59d chemosensory neuron (blue dots).

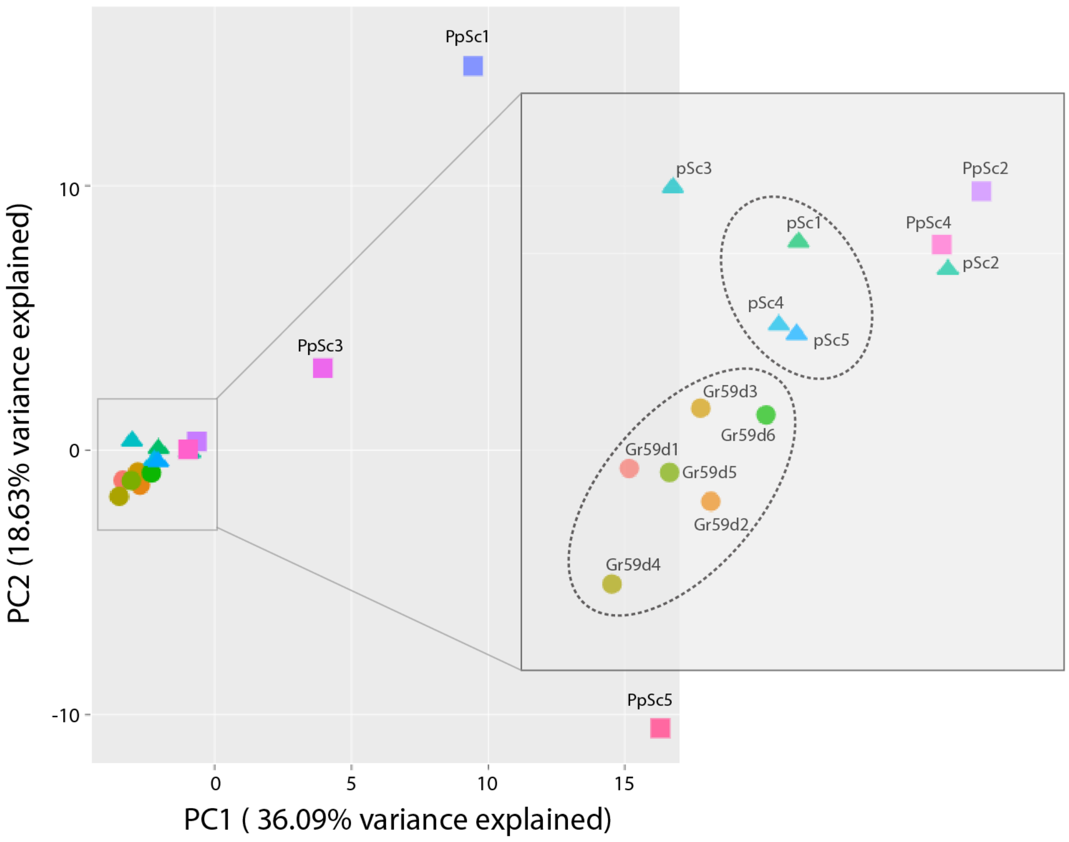


**Supplemental Figure 5 – PCA of adult Gr59d neuron, adult pSc neuron, and pupal pSc neuron samples.** The adult Gr59 neurons clustered together, but only three of the adult pSc neuron samples clustered together, and only two of the pupal pSc neurons clustered together.

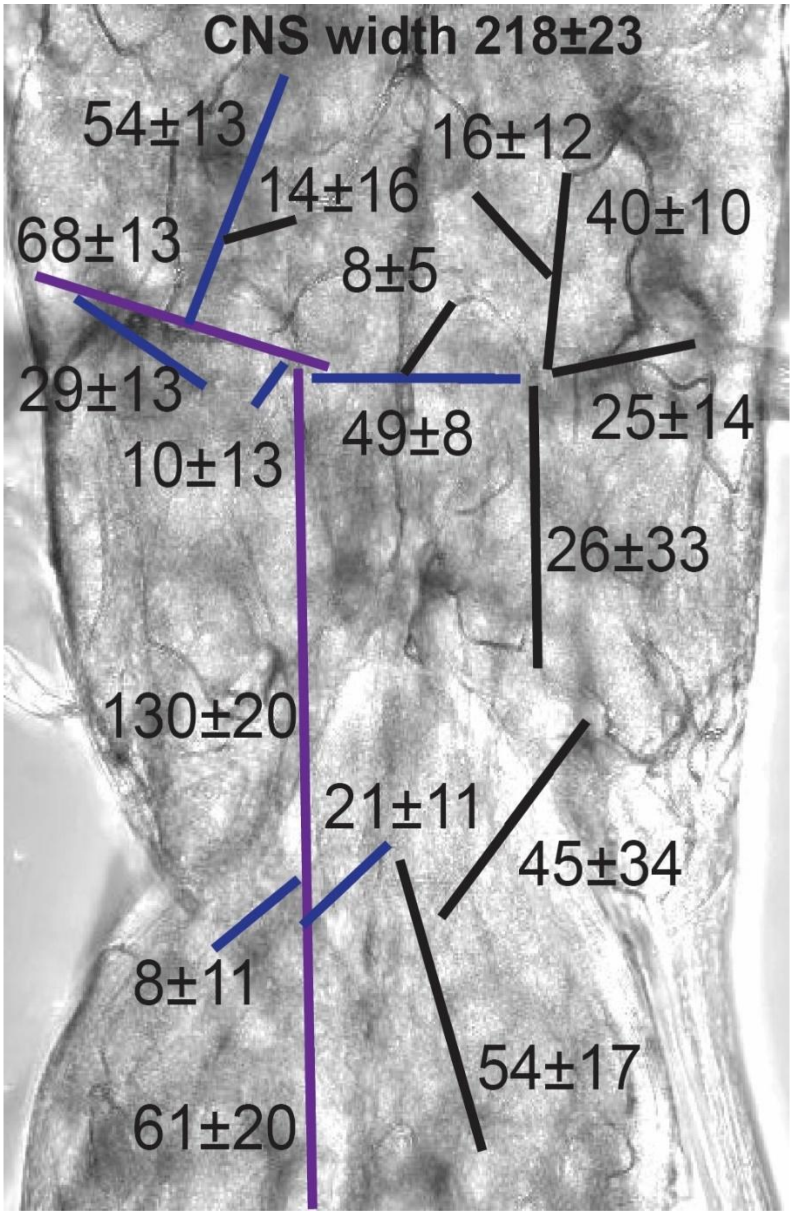


**Supplemental Figure 6 – A schematic of the wildtype pSc axonal arbor was established based on branch length and location.** Quantitative analysis of wildtype pSc axonal arbors showed 17 core branches present in over 60% of animals. 12 primary and secondary branches were observed in all animals, while two secondary branches were present in >80% of animals and three tertiary branches were found in >60%. Based on this pSc skeleton, only seven variable branches were detected in <10% of control animals. Purple lines represent primary branches; blue lines represent secondary branches; black lines represent tertiary branches. Values are expressed in µm ± S.D.

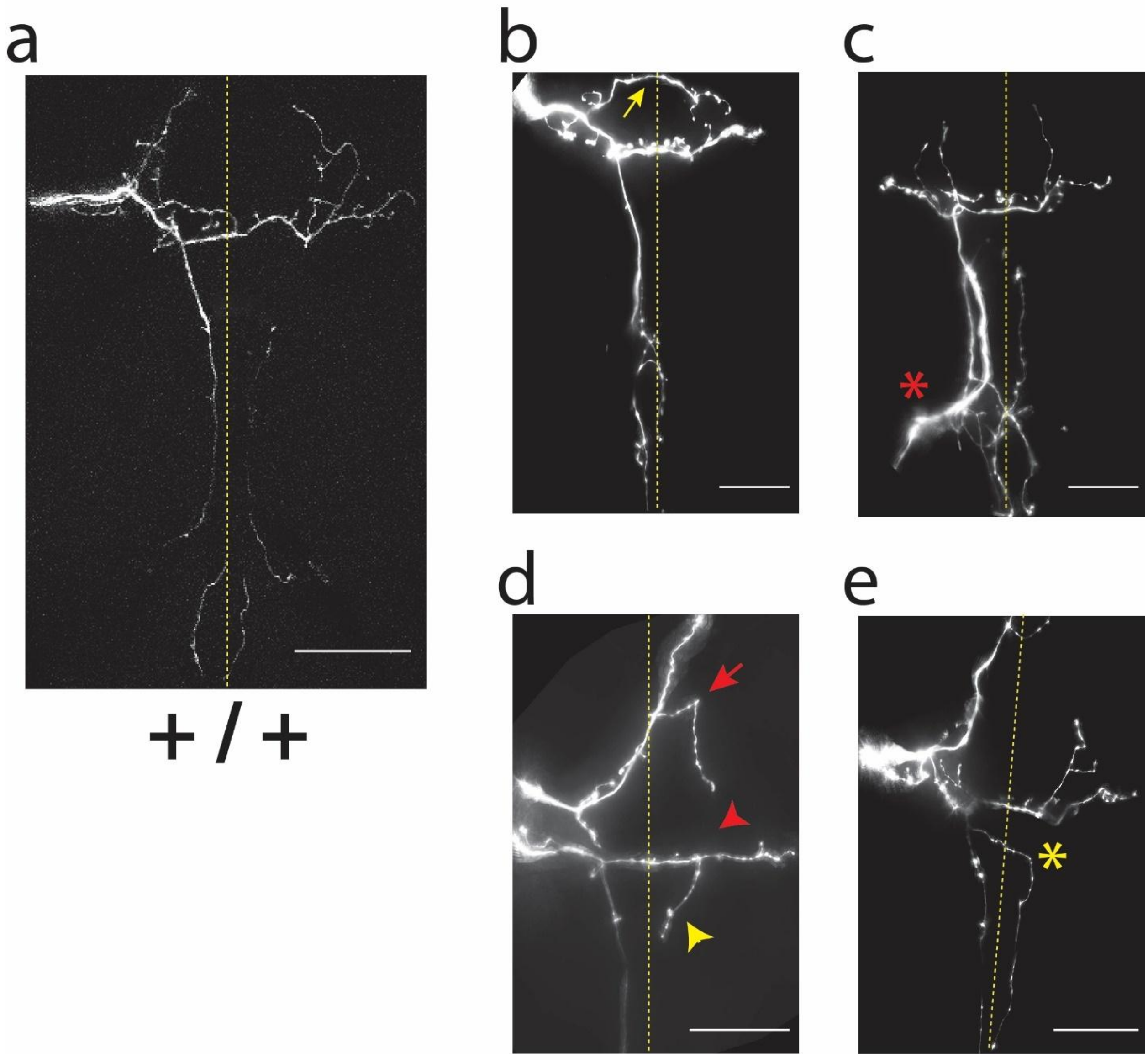


**Supplemental Figure 7 – Axonal targeting errors were divided into four categories: targeting errors, misrouting errors, guiding errors and growth errors (ectopic branches and stunted branches). a – e**, A wildtype axonal arbor and representative images of stereotyped axonal arbor errors found in RNAi knockdown mutants are shown. Red arrow points to an ectopic branch; red arrowhead points to absent branches. Yellow arrow points to targeting errors with midline crossing; yellow arrowhead points to a stunted branch. Red asterisk points to a guidance error, and yellow asterisk points to a misrouting error. The yellow dashed line represents the midline. Scale bar is 50μm.

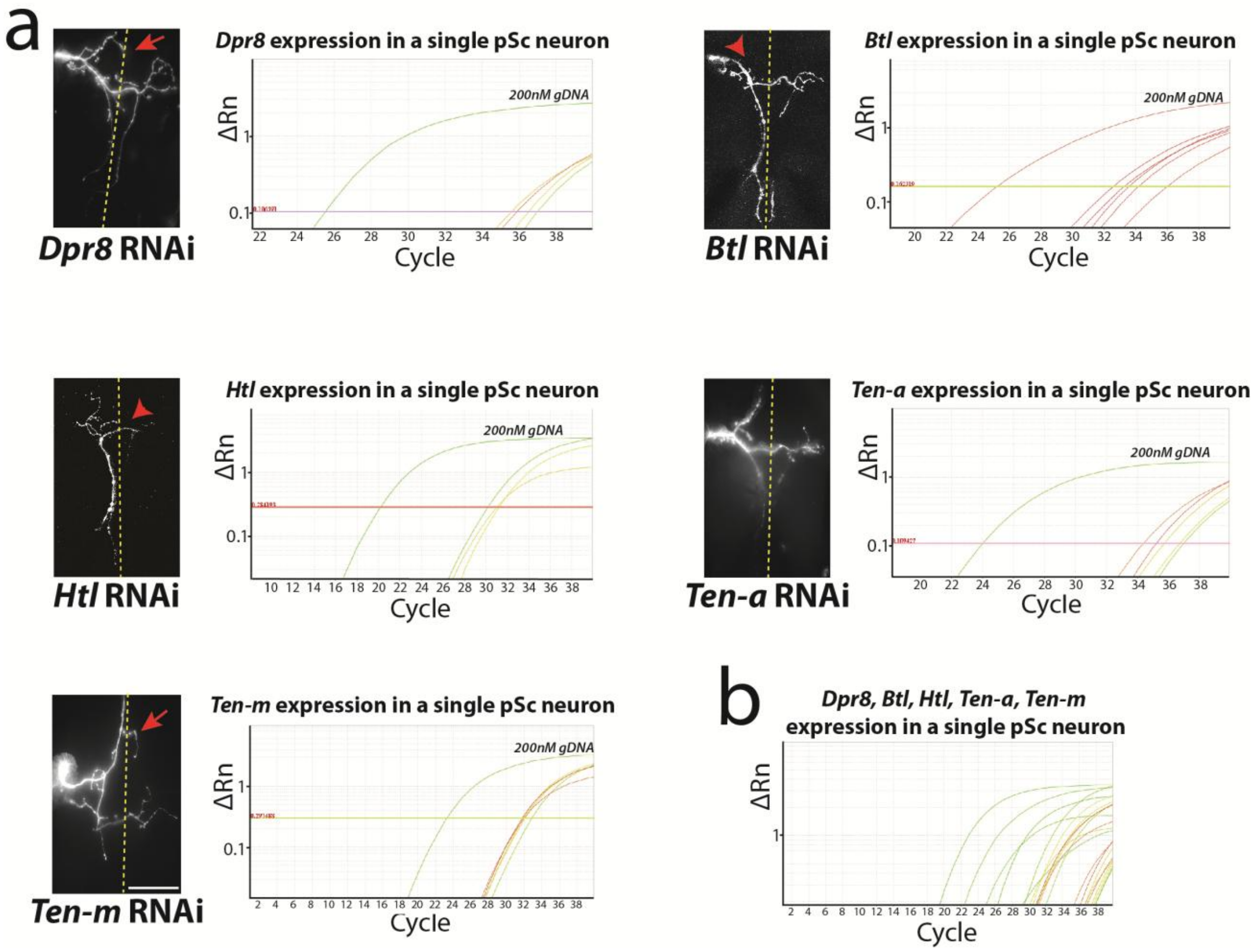


**Supplemental Figure 8 – Single neuron quantitative RT-PCR validation of the RNA sequencing data.** **a**, Quantitative reverse transcription PCR amplification curves from single wildtype pSc neurons are shown for five sample gene targets *Dpr-8, Btl, Htl, Teneurin-A* and *Teneurin-M* with its 200nM genomic DNA as a standard curve for comparison. A representative example of the RNAi knockdown for each gene is shown to the left of each gene's curve. Threshold values are shown in red numerals above its threshold line. Red arrows point to ectopic branches; red arrowheads point to missing branches. Dashed line represents the midline; scale bar is 50µm.The heatmap illustrates the differences in the expression among the *Dpr-8, Btl, Htl,* genes between seven pSc sequencing libraries. **b**, The endogenous *Dpr-8, Btl, Htl, Ten-a* and *Ten-m* expression is shown.

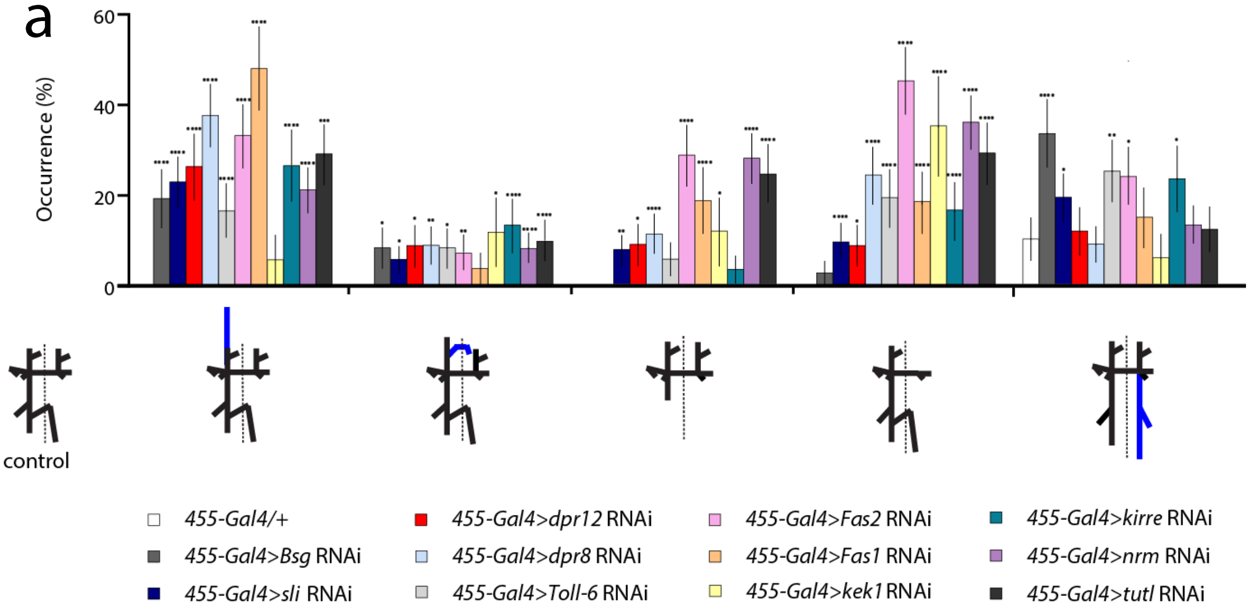


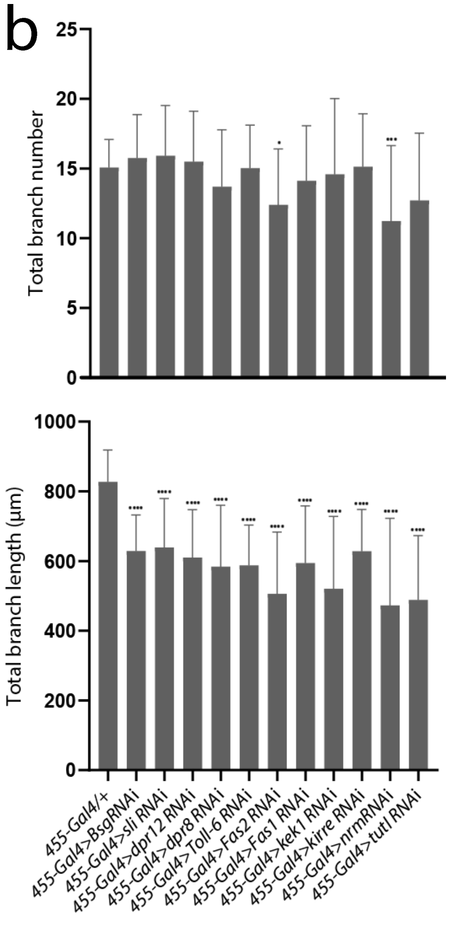


**Supplemental Figure 9 – Quantitative analysis of RNAi of differentially expressed gene within the pSc neuron.** **a,** RNAi of genes that are significantly higher expressed in pSc mechanosensory neurons than in the Gr59d neuron resulted in axonal targeting errors. Five types of axonal targeting errors were identified, shown under the x-axis in schematic. Blue branches indicate ectopic branches. The frequency of occurrence for each error type is compared to *455-Gal4 / +* controls. Error bars are standard error of the mean. **b,** RNAi of differentially expressed genes reduced pSc axonal arbour complexity. Total branch numbers and lengths for each RNAi experiment compared to control are shown. All RNAi genes had significantly reduced total branch lengths. Knockdown of *Fas2* and *nrm* resulted in a significantly reduced total branch number as well. Error bars are standard deviations. Statistical significance comparisons to control are indicated directly above the bar representing each genotype. No asterisks indicate the comparison was not significant. * $p < 0.05$, ** $p < 0.01$, *** $p < 0.001$, **** $p < 0.0001$.

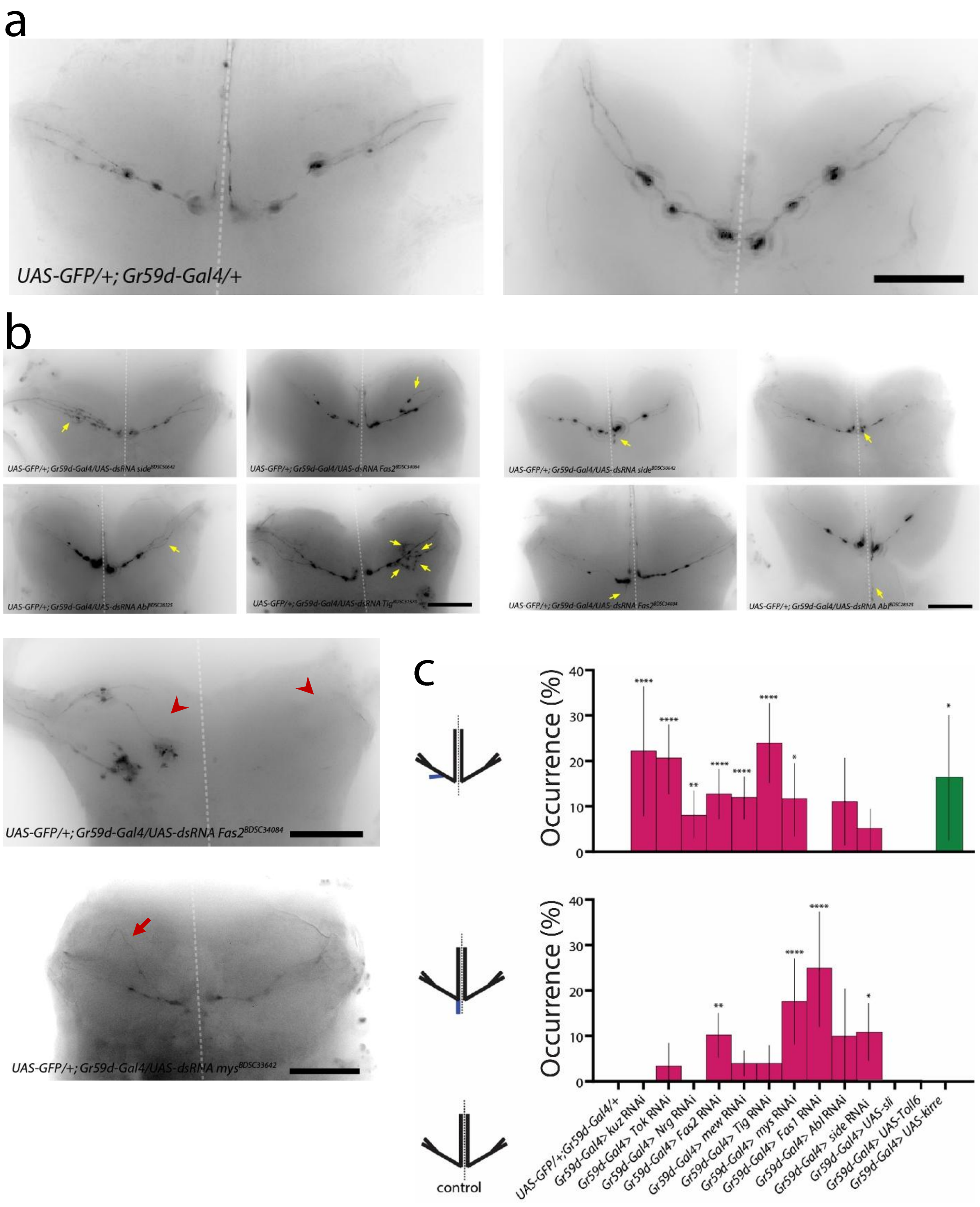


**Supplemental Figure 10 – Manipulating levels of expression of differentially expressed genes in the Gr59d chemosensory neuron resulted in axonal targeting errors. a,** Representative images of the wildtype Gr59d axons in *Gr59d-Gal4* > *UAS-GFP* animals. The Gr59d axon produces an invariant arbour within the thoracic ganglion, entering through the prothoracic neuromere and exiting without touching the midline. **b,** Representative images of

RNAi knockdown of different differentially expressed cell surface receptor genes. Various axonal targeting errors were observed, including extra branches forming before reaching the midline (yellow arrows), ectopic branch formation at the midline (yellow arrows), stalling on one side (red arrowhead) and failure to enter on the other side (red arrowhead), and premature defasciculations (red arrow). The dotted line is the midline, scale bars are 50µm. **c,** The frequency of occurrence for two targeting errors types are shown. The error types are shown as schematics to the left of the y-axis, where blue branches indicate ectopic branches. Significantly more ectopic branches were observed when *kuz*, *tok*, *Nrg*, *Fas2*, *mew*, *Tig*, and *mys* are knocked down in the Gr59d neuron. Overexpression of *kirre* in the Gr59d neuron also resulted in significantly more ectopic branches near the axonal entry point. An ectopic branch posterior extension along the midline occurred significantly more frequently when *Fas2*, *mys,* and *Fas1* were knocked down. Overexpression of *sli* or *Toll-6* alone did not result in any of these two branching errors. Magenta columns are genes knocked down using RNAi (i.e., significantly higher expressed in the Gr59d chemosensory neuron in single cell RNA sequencing). Green columns are genes that are overexpressed using UAS (i.e., significantly higher expressed in the pSc mechanosensory neuron. Statistical significance comparisons to control are indicated directly above the bar for each genotype. No asterisks indicate the comparison was not significant. Statistical significance was determined using a two-tailed *t* test for proportions. * $p < 0.05$, ** $p < 0.01$, *** $p < 0.001$, **** $p < 0.0001$.

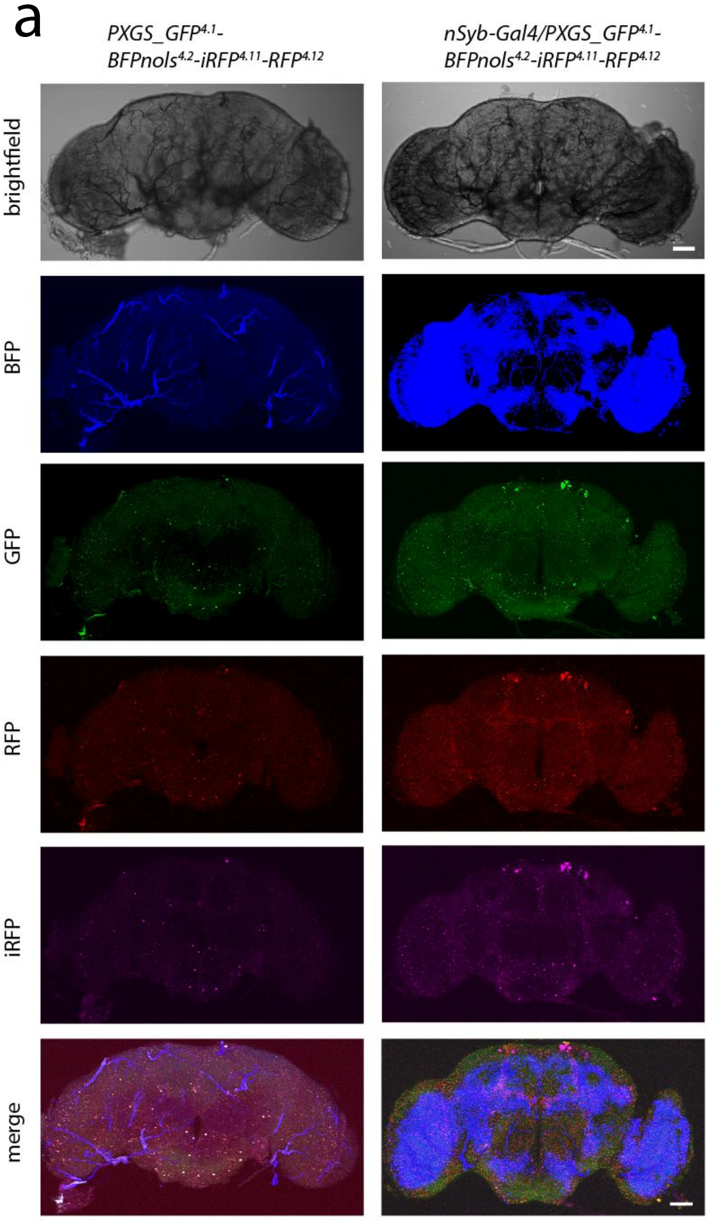


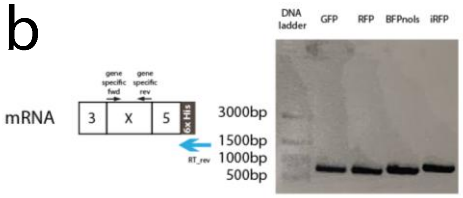


**Supplemental Figure 11 – PXGS can express multiple fluorophores simultaneously *in vivo*.** **a,** Exon 4.1, 4.2, 4.11, and 4.12 are replaced with mNeonGreen, $BFP_{nols}$, iRFP, and RFP, respectively, in the PXGS. *nSyb-Gal4* is used to drive expression of UAS-PXGS_$GFP^{4.1}$-$BFP_{nols}{}^{4.2}$-$iRFP^{4.11}$-$RFP^{4.12}$ *in vivo*. The brains of flies without a Gal4 driver as a control are shown on the left. In the brains of experimental animals (right images), the $BFP_{nols}$ signal was observed as obvious puncta throughout the brain, but the cytoplasmic GFP, RFP, and iRFP signals were less obvious without the subcellular localization signals that were included in later versions. The scale bar is 50µm. **b,** RT-PCR of each fluorophore was performed on brain extracts to verify that all transgenes were spliced properly.

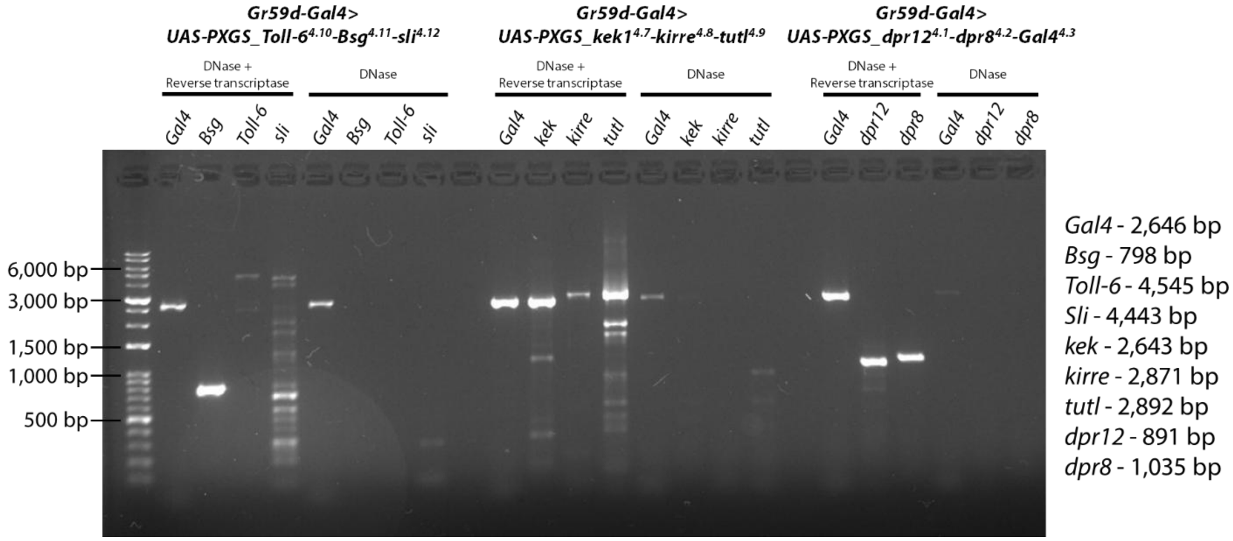


**Supplemental Figure 12 – Gr59d-Gal4 can express cell surface receptor genes *in vivo* using PXGS.** The Gr59d chemosensory neuron was dissected from animals expressing different cell surface receptor genes using PXGS (*Gr59d-Gal4 > UAS-PXGS_Toll-6*$^{4.10}$*-Bsg*$^{4.11}$*-sli*$^{4.12}$; *Gr59d-Gal4 > UAS-PXGS_kek1*$^{4.7}$*-kirre*$^{4.8}$*-tutl*$^{4.9}$; and *Gr59d-Gal4 > UAS-PXGS_dpr12*$^{4.1}$*-dpr8*$^{4.2}$*-Gal4*$^{4.3}$), and RT-PCR was performed. The *6×His* genetic/RNA tag was used for the PXGS-specific reverse primer. Specific primers for each transgene were used for PCR, with the expected PCR band sizes shown on the right.

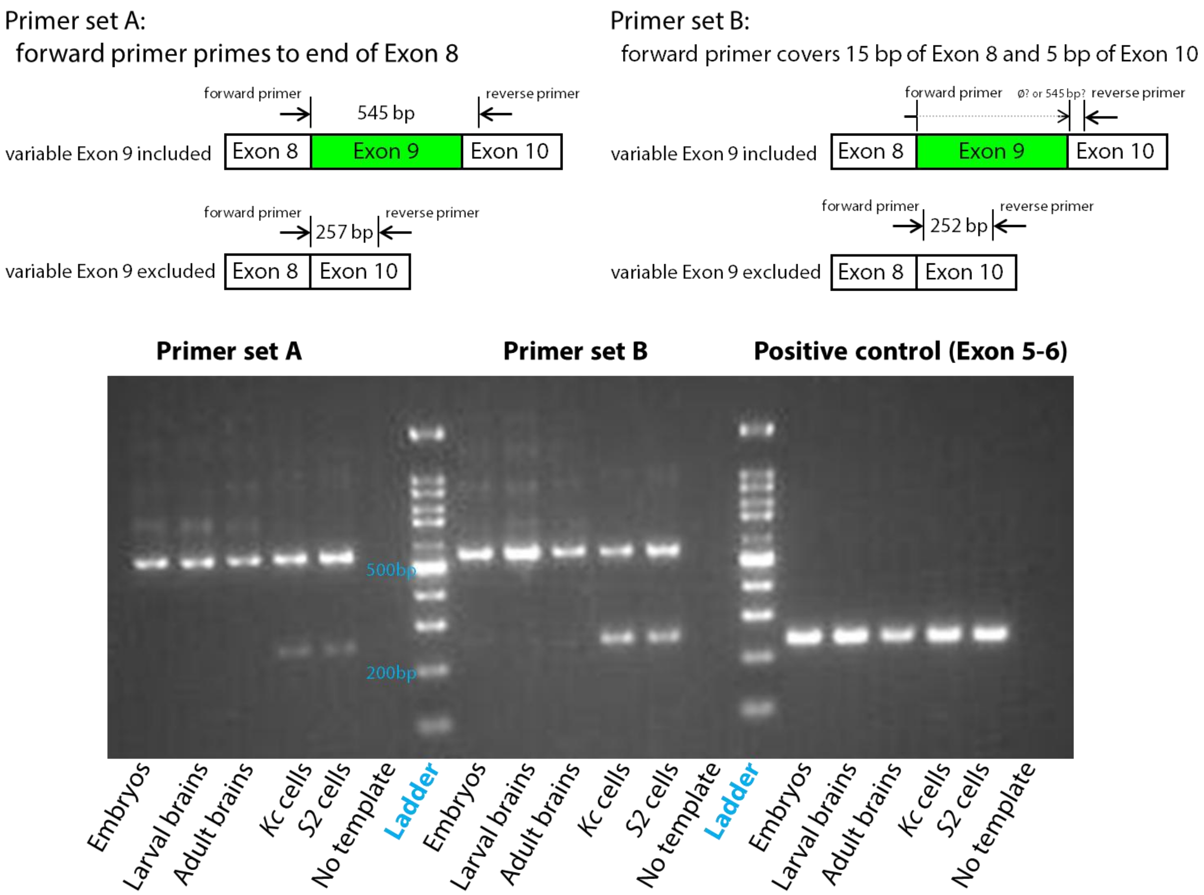


**Supplemental Figure 13 – *Dscam* variable exon 9 is skipped in *Drosophila* cell lines.** To verify the skipping of variable exon 9 in *Kc* cells from the single cell long read-length single molecule real-time RNA sequencing results, we performed reverse transcription PCR on different *Drosophila* tissues and cell lines. We found that variable exon 9 was skipped in *S2* cells and *Kc* cells.
